# Mapping the Changing Neural Architecture of Narrative Processing Using Naturalistic Stimuli: an fMRI Study

**DOI:** 10.1101/2023.09.15.557976

**Authors:** Christina Haines, KiAnna Sullivan, Keva Klamer, Joshua Craig, Chelsea Ekstrand

**Affiliations:** Ekstrand Neuroimaging Lab, Department of Neuroscience, University of Lethbridge, 4401 University Dr W, Lethbridge, AB, Canada, T1K 6T5

## Abstract

A narrative is a coherent representation of actual or fictional events designed to connect experiences. Narratives provide a unique opportunity to investigate brain functions in scenarios more closely resembling real-world experiences. However, most neuroimaging studies examining narrative formation have utilized static stimuli that fail to capture the intricacies of narrative construction in everyday life, particularly how cognitive demands change over the course of narrative processing. The current research uses functional magnetic resonance imaging (fMRI) to examine dynamic narrative processing over the course of a full-length audiovisual narrative. We examined changes in neural synchrony (as quantified by intersubject correlations) in areas related to semantic memory, episodic memory, and visuospatial attention between the beginning, middle, and end of the narrative. Results from two experiments identified two core narrative processing networks responsible for constructing coherent representations across extended timescales. The first network is associated with the early narrative construction, and includes the right intraparietal sulcus/superior parietal lobule, bilateral angular gyrus, bilateral precuneus, and left fusiform gyrus. The second network consists of the right ventral frontal cortex and bilateral parahippocampal cortices, and is associated with longer term narrative integration. Together, these regions provide the framework for successful narrative processing during naturalistic stimuli.

## Neural Correlates of Narrative Structure During Naturalistic Audiovisual Film Using Functional Magnetic Resonance Imaging

A narrative is a coherent representation of actual or fictional events designed to connect experiences (Martinez-Conde et al., 2019). Narrative comprehension is complex, requiring sustained attention and integration of cognitive inputs. Narratives can come in various forms, including written word, spoken stories, film, video games, and virtual reality (Willems et al., 2020). The encoding of narratives relies on an innate human ability to process semantic information, recall memories, engage with emotional content, and interact socially. Through this intricate cognitive process, individuals can perceive, categorize, construct, and store a cohesive representation of information. As new events and experiences are encountered, the brain can adapt and transform mental representations of people, events, and environments to align with the dynamically changing world. Not only can these representations be modified, but they can also be assigned to a mental timeline, creating a sense of temporal coherence. Further, the presence of new information can prompt individuals to revise and update existing narrative representations, allowing for a flexible and adaptive understanding of personal stories. Thus, narratives serve as a means for connecting experiences and constructing meaningful representations of the world.

### Cognitive Processes Required for Narrative Processing

Narrative formation is complex and engages various short-term and long-term cognitive processes to comprehend and create meaningful representations. First, visuospatial attention is involved in directing cognitive resources toward specific aspects of the narrative, such as characters, events, or plot developments (Jääskeläinen et al., 2020). The visuospatial attention network consists of two right-lateralized cortico-cortical neural networks (Corbetta & Shulman, 2002), a dorsal attention network responsible for goal-directed attention (frontal eye field (FEF) and intraparietal sulcus (IPS)/superior parietal lobule (SPL)) and a ventral attention network involved in bottom-up attentional selection (temporoparietal junction (TPJ) and ventral frontal cortex (including the inferior frontal gyrus (IFG) and middle frontal gyrus (MFG)). Second, semantic memory integrates knowledge about the world, including concepts, facts, and general knowledge, that individuals have accumulated over their lifetime (Binder et al., 2009). The semantic system is primarily left-lateralized and includes the temporal pole, anterior cingulate cortex (ACC), fusiform gyrus, intraparietal sulcus (IPS), posterior middle temporal gyrus (MTG), and prefrontal cortex (PFC). Third, episodic memory is responsible for encoding and retrieving personal experiences or episodes (Ritchey & Cooper, 2020), and plays a crucial role in recollecting and reconstructing memory representations built throughout exposure to the narrative. The posterior medial episodic network provides a neural basis for the construction of dynamic and detailed mental representations and consists of the hippocampus, parahippocampal cortex (PHC), retrosplenial cortex (RSC), precuneus, angular gyrus (AG), posterior cingulate cortex (PCC), and medial prefrontal cortex (mPFC) (Ritchey & Cooper, 2020). Overall, visuospatial attention, semantic memory, and episodic memory are interconnected cognitive processes that contribute to the comprehension and encoding of narratives and these processes rely on neural networks distributed across various brain regions (see Figure 1).

**Figure 1.**
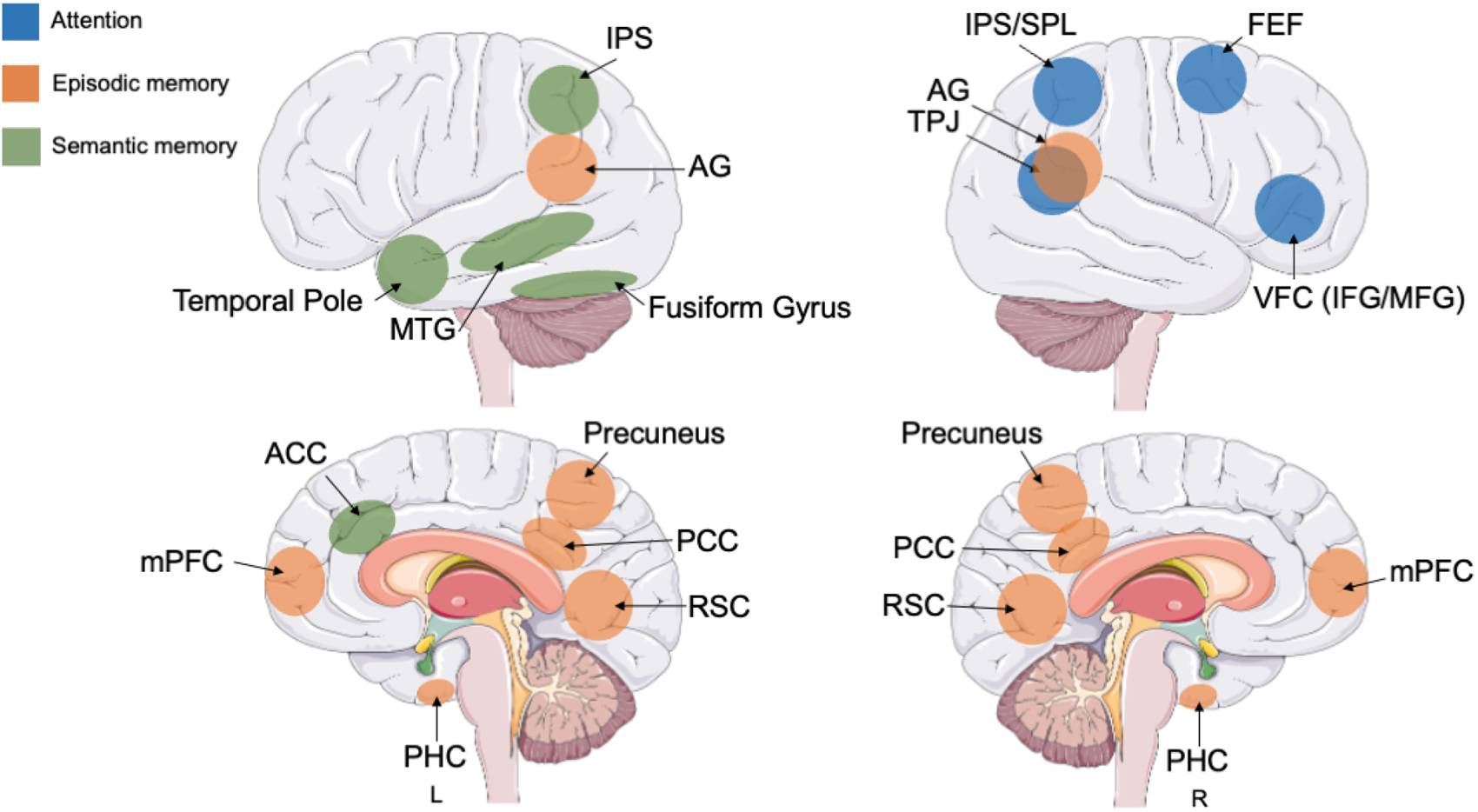
Narrative Processing Model. This figure depicts the regions we hypothesize to be involved in narrative processing. Three prominent cognitive systems—semantic memory, episodic memory, and attention—are shown with their respective neural correlates. Brain model downloaded from Servier Medical Art (“SMART – Servier Medical ART,” n.d.) licensed under a Creative Commons Attribution 3.0 unported license.

### Naturalistic Cognitive Processing

Exploring how narratives are integrated in the brain within experimental settings presents a set of challenges. Two major obstacles are that narrative processing is complex and multimodal, and processing unfolds across extended timescales—ranging from minutes to years—making it difficult to capture effectively in conventional research designs. To deal with these complexities, an emerging consensus proposes integrating naturalistic stimuli into research protocols, which better replicate the perceptual, cognitive, and emotional demands inherent in real-life (Vanderwal et al., 2019). Examples of naturalistic stimuli include audiovisual movie clips, TV advertisements, spoken narratives, interactive encounters with virtual agents, gaming environments, and virtual reality (Sonkusare et al., 2019). However, with this shift to naturalistic stimuli comes the computational challenge of analyzing and interpreting complex neural responses associated with multiple task demands. To address this, intersubject correlation (ISC) analysis has been proposed (Hasson et al., 2010; Nastase et al., 2019). ISC analysis identifies neural activity that is shared between subjects by examining correlations in hemodynamic responses across the time course of a naturalistic stimulus (Pajula et al., 2012). The primary objective of ISC analysis is to uncover shared neural activity, also known as neural synchrony, among a large proportion of participants. Using this analysis, it is possible to create maps illustrating the distribution of neural synchrony across the brain. ISC analysis is commonly employed with naturalistic stimuli as it can reveal networks of brain regions that consistently synchronize during certain tasks, experiences, or cognitive processes, providing insight into how humans process and react to the same or similar situations (Hasson et al., 2010).

Previous research on the neural mechanisms of narrative understanding has consistently identified several brain areas, including the precuneus, PCC, TPJ, and temporal pole (Cavanna & Trimble, 2006; Utevsky et al., 2014). For example, previous research from Tylén et al. (2015) used fMRI and incoherent and coherent podcast episodes to study the neural correlates of plot formation. A two-level general linear model (GLM) analysis contrasting coherent events to incoherent events showed significant activation in the PCC, precuneus, anterior prefrontal cortex, orbitofrontal cortex, right hippocampus, and right caudate nucleus, as well as large areas of the temporal lobes extending from the temporal pole to the TPJ. Kauttonen et al. (2018) looked at the formation of key plot points of a narrative in the brain over longer timescales using naturalistic stimuli. To do so, 25 participants watched one of two versions of the film *Memento* (Nolan, 2001) during fMRI, one presented in reverse chronological order (the original version of the film) with brief scenes that overlap and serve as memory cues for the viewer, and a version of the film edited to follow chronological order. Multivariate eventlJrelated pattern analysis and representational similarity analysis revealed significant clusters of brain activity in the frontal pole, cingulate and paracingulate gyrus, precuneus, AG, and the MFG when comparing between participants who watched the film in reverse chronological order versus those who watched it in chronological order. These results suggest the role of these regions in coherent plot formation.

Overall, these studies provide valuable insight into the neural correlates of narrative processing. Thus, based on previous research, the areas proposed to be involved in narrative processing in naturalistic stimuli include attentional areas such as the right FEF, SPL, IFG/MFG, and TPJ, semantic processing areas including the left temporal pole (hub), ACC (hub), fusiform gyrus, IPS, MTG, and PFC, and episodic memory areas such as the mPFC, precuneus, PHC, hippocampus, RSC, AG, PCC (see Figure 1). However, one aspect that remains largely unexplored is the dynamic nature of how cognitive demands evolve over the course of a narrative. While existing research has focused on specific cognitive processes such as cued recall, the role of context in narrative, and plot formation on a short time scale, there is a need to investigate how these processes interact and transform as the narrative unfolds. By examining evolving cognitive demands within a narrative, we can gain a more comprehensive understanding of the complex interplay between cognitive processes and the narrative structure, and how individuals adapt their cognitive resources to meet the changing requirements of long-term memory integration.

### Classical Film Plot Development

Films provide extensive multimodal and emotionally laced information to viewers, which provides an ideal model to assess the way the brain processes real-world information in a research setting. They also allow for the assessment of narrative processing on a more defined scale, rather than in the expansive sense of one’s own narrative. Based on previous research and drawing on classic film theory, it has been established that the structure of a “classic Hollywood film” consists of three acts: Set-up, Development, and Resolution (Cutting, 2016). This continues to be widely employed in present-day Hollywood movies (Brütsch, 2015; Guha et al., 2015). On average, Set-up is the first third of the narrative, Development occupies the middle third, and Resolution forms the final third. Act I, known as the Set-up act, introduces the protagonist and establishes their world and overarching goals. Act II, the Development act, primarily revolves around the protagonist facing obstacles. The main character typically reaches a pivotal moment where they must decide whether to continue pursuing the goal or give up. Act III, the Resolution act, centers on the protagonist’s ability to overcome the ultimate obstacle, leading to the climactic point and a conclusion.

### Current Research

This research sought to examine the brain regions involved in narrative processing over time using a naturalistic paradigm. To achieve this, we used data from the Naturalistic Neuroimaging Database (NNdB; version 2.0.0; Aliko et al., 2020) of participants who watched a full-length audiovisual movie during fMRI. This film was divided into sections based on the classical film plot theory previously discussed: Set-up, Development, and Resolution. By examining each section individually and analyzing the differences in neural synchrony between the stages, this research aimed to unravel changes in brain synchrony associated with changing cognitive task demands as the storyline progressed. Our general hypothesis was that, over the course of the narrative, cognitive demands would change, thus leading to changes in neural synchrony in attention, semantic processing, episodic memory, and narrative processing regions across the three acts.

## Experiment 1

### Introduction

In Experiment 1, our objective was to investigate neural synchrony in regions linked to episodic memory, semantic memory, attention, and overall narrative processing throughout distinct stages of a complex, audiovisual narrative. To achieve this, we used fMRI data from participants who watched the film *500 Days of Summer* (Webb, 2009). This film was chosen due to its adherence to the three-act structure commonly employed in contemporary Hollywood movies (Sharma & Rajamanickam, 2015). We divided fMRI data into three equal segments based on the total number of volumes acquired, which correspond to the Set-up, Development, and Resolution acts. It is important to note that our division of the data was designed to capture potential changes in cognitive processes across the progression of the narrative. In general, we hypothesized that synchrony in brain areas associated with attention, semantic memory, episodic memory, and narrative processing would be unique for the Set-up, Development, and Resolution phases. Specific hypotheses for each section are outlined below.

During the Set-up phase, focused on world-building and memory formation, we hypothesized that the right FEF, IPS/SPL (goal-directed visuospatial attention), left ACC, temporal pole, IPS, fusiform gyrus, MTG (semantic memory), bilateral AG, precuneus, PCC, RSC, PHC, hippocampus, and mPFC (episodic memory) would exhibit significant synchrony. In the Development phase, where characters face obstacles and make decisions, we expected decreased synchrony in the right FEF and IPS/SPL (goal-directed visuospatial attention) but continued synchrony in bilateral AG, precuneus, PCC, RSC, PHC, hippocampus, mPFC (episodic memory), left ACC, temporal pole, IPS, fusiform gyrus, MTG (semantic memory), right TPJ and IFG/MFG (stimulus-driven visuospatial attention). In the Resolution phase, tied to drawing inferences and the story’s conclusion, we predicted decreased synchrony in the right FEF, IPS/SPL (goal-directed visuospatial attention), left ACC, temporal pole, IPS, fusiform gyrus, MTG (semantic memory), bilateral AG, precuneus, PCC, RSC, PHC, hippocampus, and mPFC (episodic memory), while the right TPJ and IFG/MFG (stimulus-driven visuospatial attention) would show synchronous activity due to the absence of new plot-driven information.

### Methods

#### Participants

For this study, we used preprocessed fMRI data from the NNdB (Aliko et al., 2020) v2.0 of twenty participants (10 female, average age = 27.7 years) who watched the film *500 Days of Summer* (Webb, 2009) during fMRI. *500 Days of Summer* (Webb, 2009) was selected from the 10 films utilized in the NNdB for the following reasons: This film follows the classic 3-act structure and has the largest number of acquired datasets per movie in this database. All participants were right-handed, native English speakers, had unimpaired or corrected-to-normal vision, no hearing impairments, and had no history of neurological or psychiatric illnesses (see Aliko et al., 2021 for detailed information).

#### Data Acquisition

All data acquisition details can be found in Aliko et al. (2021). In summary, MRI data was collected on a 1.5T Siemens Magnetom Avanto scanner with a 32-channel head coil. FMRI data was acquired using an echo planar imaging (EPI) sequence with a 4x multiband factor and the following scanning parameters: repetition time (TR) of 1s, echo time (TE) of 54.8ms, flip angle of 75°, and a resolution of 3.2mm isotropic. A T1-Magnetization Prepared Rapid Acquisition Gradient Echo anatomical scan was acquired with a TR of 2.73s, TE of 3.57ms, and a resolution of 1.0mm³. To ensure optimal audio quality, noise-attenuating headphones were used. Participant attention was closely monitored using a camera fixated on their eyes. The visual part of the film was presented through a mirror-reversing LCD projector. To maintain the naturalistic viewing experience and minimize interruptions, the film was played with minimal breaks. However, due to the limitations of the EPI sequences and software, the film was played in 40–50-minute segments. These breaks were intentionally timed to occur during scenes that did not contain crucial plot information or dialogue.

#### Data Preprocessing

Data preprocessing was conducted by Aliko et al. (2021). Briefly, because the films were obtained in separate runs to accommodate EPI and software requirements, the time series were concatenated after undergoing timing correction preprocessing. In addition to the concatenation, all functional scans underwent the following preprocessing steps: time shifting, despiking, volume registration, MNI alignment, mask time-series, smoothing 6mm FWHM, detrending with regressors (e.g., for motion), timing correction, manual ICA denoising (Aliko et al., 2020) using AFNI’s afni_proc.py pipeline (Chen et al., 2017; Cox, 1996). For this analysis, we utilized preprocessed functional files that had undergone blurring and remained uncensored. We divided each functional data file into three sections: Set-up, Development, and Resolution, using *fslroi* (Jenkinson et al., 2012). For *500 Days of Summer* (Webb, 2009) the film consisted of 5470 acquired volumes. The Set-up section of the film spanned volumes 1 – 1823, the Development section spanned volumes 1824 – 3647, and the Resolution section consisted of volumes 3648 – 5470. Because 5470 is not divisible by three, the Resolution section of the film consisted of one less volume than the Development and Resolution sections.

#### Data Analysis

Using the data files split into the Set-up, Development, and Resolution sections, we then calculated ISCs for each pair of subjects in the analysis. These pairings were formed across 20 participants within each film section (Set-up, Development, and Resolution), resulting in 190 unique pairings of participants per section. AFNI’s *3dtcorrelate* (Cox, 1996) was used to generate these ISC pairings for each film section.

Next, we used linear mixed effects (LME) modelling implemented via the *3dISC* module in AFNI (Chen et al., 2017; Cox, 1996) to identify significant regions of neural synchrony, as quantified using ISC analysis. To do so, we ran an LME analysis for all 190 participant ISC pairings for each of the three film sections individually, resulting in three separate one-group analyses (Set-up, Development, and Resolution). LME modelling is a parametric method that assumes certain distributions of the data, such as normal or Gaussian distributions. This approach accounts for the complex covariance structure of ISC data and incorporates both random effects (e.g., variability in brain activity across participants) and fixed effects (e.g., effects of experimental conditions on brain activity) within the same model (Chen et al., 2017). This enables a more accurate estimation of fixed effects by accounting for data variability due to random effects (Koerner & Zhang, 2017). Significant ISC was determined using a voxel-wise FDR threshold of *q* < .0001 for each contrast. The critical *t* values were 5.5987, 5.6036, and 5.8193 for Set-up, Development, and Resolution, respectively. Using these critical *t*-statistics, the resulting *t*-statistic image was thresholded using *fslmaths* ‘-thr’ function (Jenkinson et al., 2012), at the aforementioned critical *t* value and the ISC image was masked using *fslmaths* ‘-mas’ function (Jenkinson et al., 2012).

### Results

#### Set-up

Results from the one-way LME for the Set-up contrast found 33 significant clusters of neural synchrony across the brain (see Figure 2 and Supplementary Materials Appendix A). In line with our hypotheses, we found significant neural synchrony in regions associated with visuospatial attention, including the right FEF, SPL, TPJ, and IFG/MFG. Further, we found significant neural synchrony in regions associated with episodic memory (bilateral precuneus, PHC, RSC, AG, PCC, and mPFC) and semantic memory (left temporal pole, ACC, fusiform gyrus, IPS, posterior MTG, and PFC). We also found significant clusters of neural synchrony in areas such as the bilateral precentral gyrus, bilateral frontal poles, and right temporal pole. In addition to these cortical areas, subcortical regions including the right thalamus, right amygdala, right caudate, and bilateral crus II in the cerebellum showed significant neural synchrony during this stage of the narrative.

**Figure 2.**
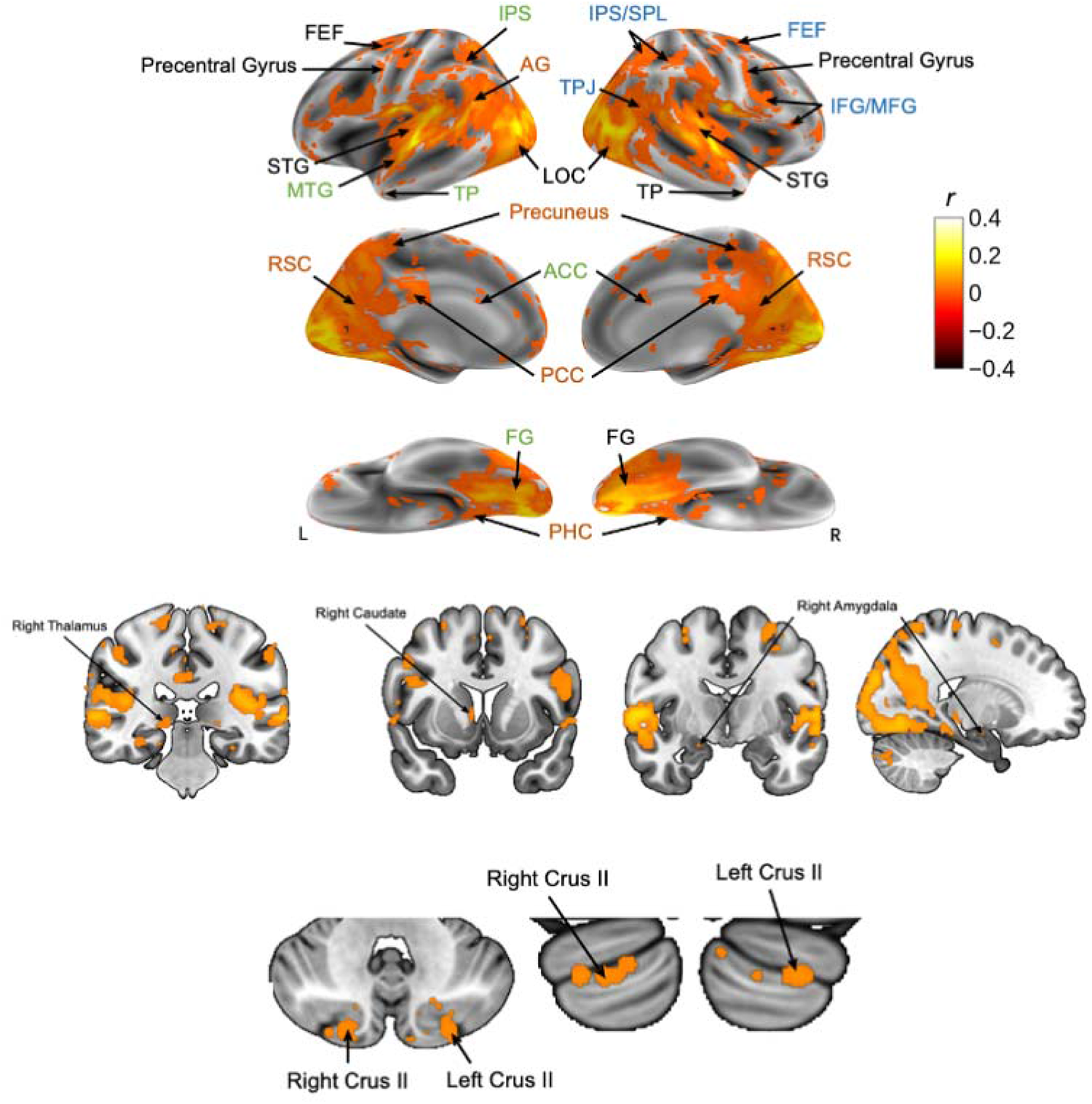
*500 Days of Summer*: Set-up Neural Synchrony. Synchrony maps derived from fMRI data during audiovisual movie watching. Maps are shown for the left (L) and right (R) hemispheres, in the lateral, medial, and inferior orientations. Areas in blue represent neural correlates of visuospatial attention, areas in green represent neural correlates of semantic memory, and areas in orange represent neural correlates of episodic memory.

#### Development

Results from the one-way LME for the Development contrast found 38 significant clusters of neural synchrony (see Figure 3 and Supplementary Materials Appendix A). In line with our hypotheses, we found significant synchrony in regions associated with visuospatial attention, namely the right FEF, SPL, TPJ, and IFG/MFG. Additionally, regions involved in episodic memory exhibited neural synchronization, particularly in the bilateral precuneus, PHC, RSC, AG, PCC, and mPFC. We found significant clusters of neural synchrony in the following semantic memory areas (left-lateralized): temporal pole, fusiform gyrus, IPS, posterior MTG, and PFC. Other areas that showed significant neural synchrony were the bilateral precentral gyrus, bilateral frontal poles, and right temporal pole, as seen in Figure 3. Further, subcortical areas with significant clusters of neural synchrony during the Development section include the right hippocampus and the bilateral crus II in the cerebellum.

**Figure 3.**
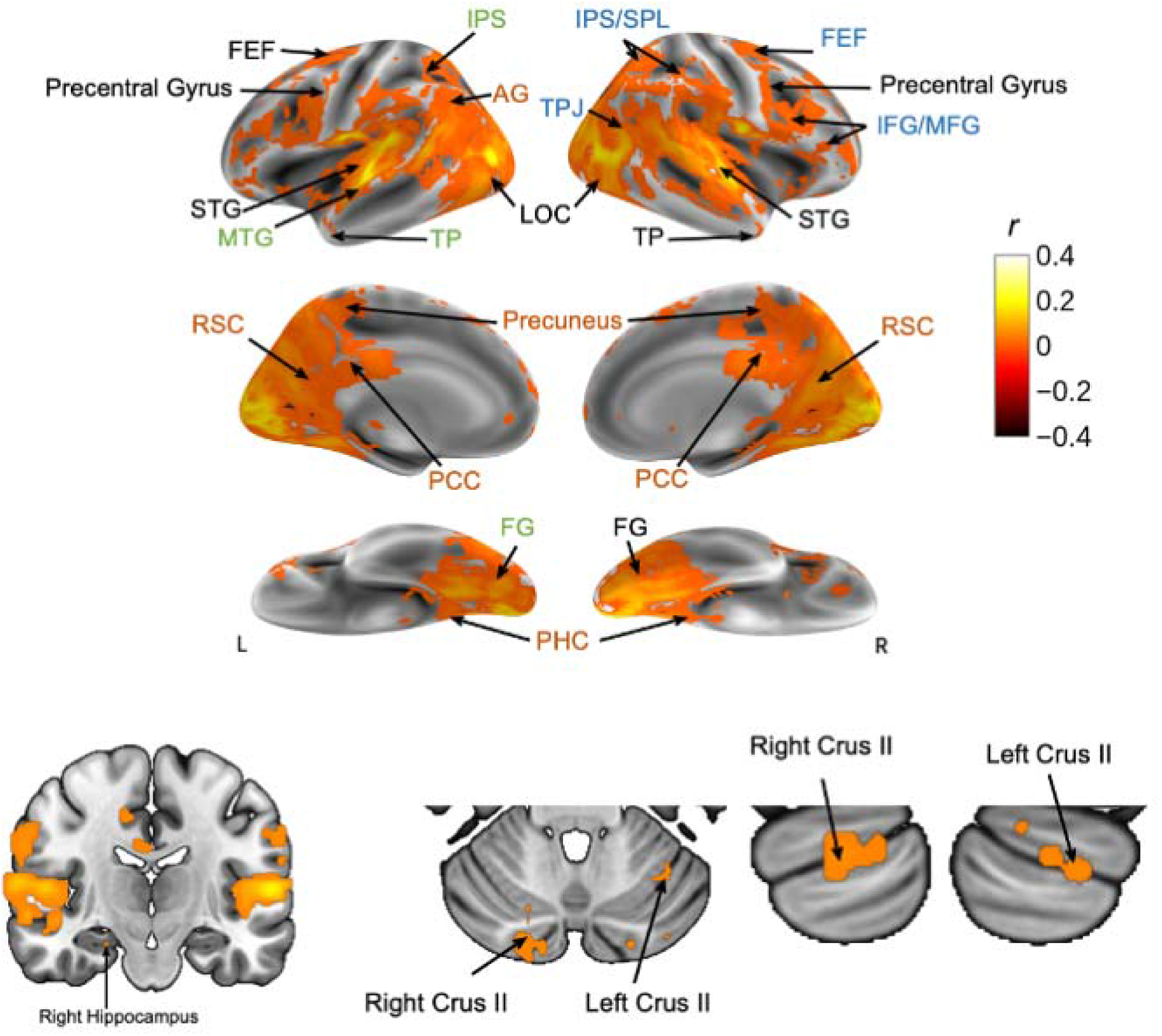
*500 Days of Summer*: Development Neural Synchrony. Synchrony maps derived from fMRI data during audiovisual movie watching. Maps are shown for the left (L) and right (R) hemispheres, in the lateral, medial, and inferior orientations. Areas in blue represent neural correlates of visuospatial attention, areas in green represent neural correlates of semantic memory, and areas in orange represent neural correlates of episodic memory.

#### Resolution

Results from the one-way LME for the Resolution contrast found 22 significant clusters of neural synchrony (see Figure 4 and Supplementary Materials Appendix A). Visuospatial attention areas, specifically the right FEF, SPL, TPJ, and IFG/MFG, showed significant synchronous activity. Moreover, regions associated with episodic memory demonstrated neural synchronization, particularly in the bilateral precuneus, PHC, RSC, AG, PCC, and mPFC. Significant semantic memory areas included the left temporal pole, fusiform gyrus, IPS, and PFC. Other areas that showed significant clusters of neural synchrony were the precentral gyrus, bilateral frontal poles, and right temporal pole, as seen in Figure 4. Subcortical areas with neural synchrony during the Resolution section include the bilateral caudate and the bilateral crus II in the cerebellum.

**Figure 4.**
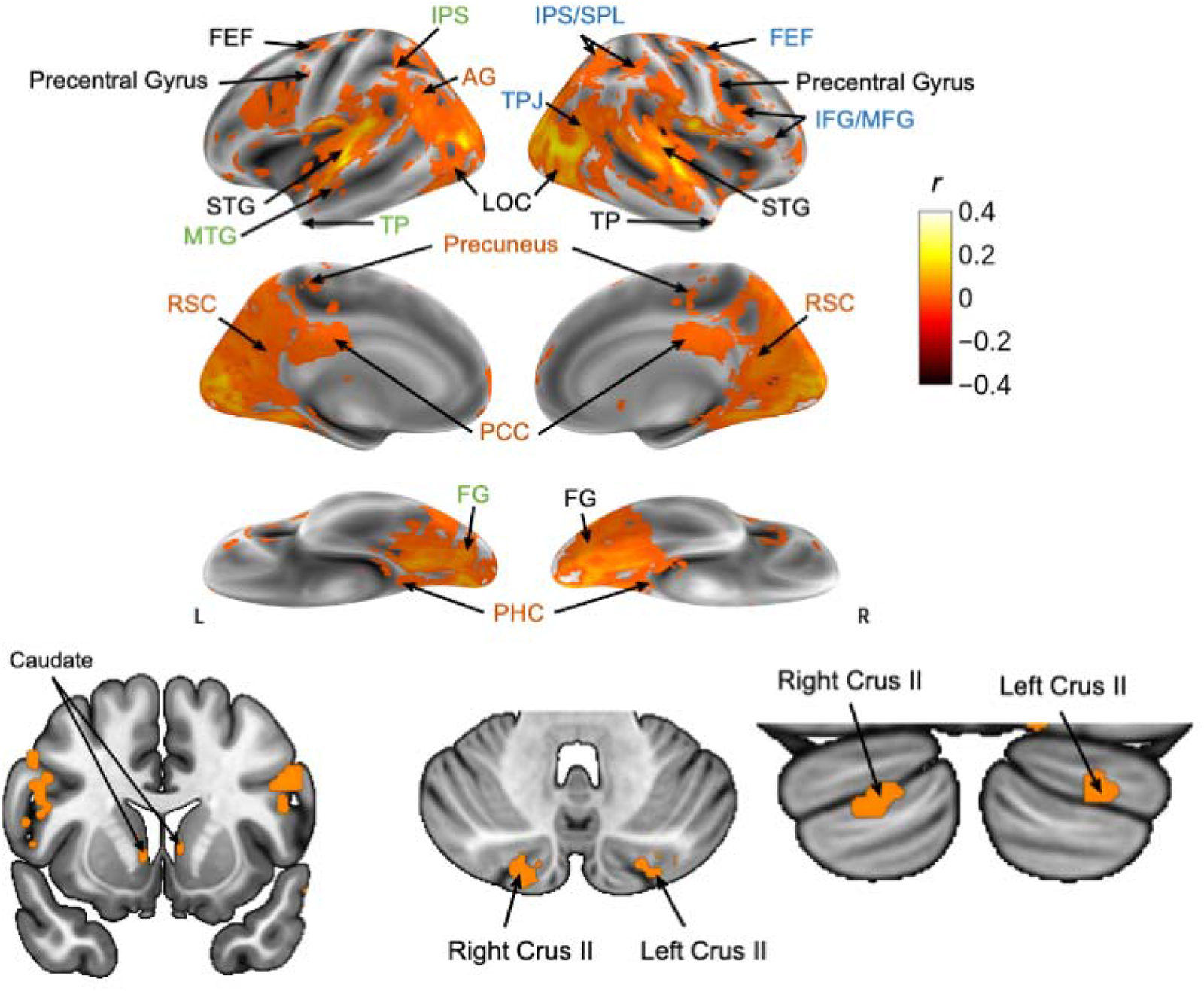
*500 Days of* Summer: Resolution Neural Synchrony. Synchrony maps derived from fMRI data during audiovisual movie watching. Maps are shown for the left (L) and right (R) hemispheres, in the lateral, medial, and inferior orientations. Areas in blue represent neural correlates of visuospatial attention, areas in green represent neural correlates of semantic memory, and areas in orange represent neural correlates of episodic memory.

### Discussion

Results from Experiment 1 provide valuable insight into the neural correlates of narrative processing, highlighting diverse cognitive processes that contribute to processing. Significant neural synchrony was observed in attentional areas, including the areas of the dorsal (right FEF and SPL) and ventral (right TPJ and IFG/MFG) attention networks across all three sections of the film. This suggests that both goal-driven attention and stimulus-driven attention are being utilized across the narrative. According to Jääskeläinen et al. (2020), the TPJ, IPS, and FEF are involved in narrative engagement and arousal. Episodic memory areas showed consistent bilateral synchrony across all three narrative sections, including the precuneus, PHC, RSC, AG, PCC, and mPFC. This suggests narrative processing requires significant episodic memory system recruitment to parse and encode important episodic narrative information. In addition, the precuneus is involved in narrative interpretation, highlighting its role in connecting narrative elements and interlacing them into a coherent representation (Jääskeläinen et al., 2020). The right hippocampus, an area known to be involved in episodic memory, showed significant neural synchrony during the Development portion of the film. This finding mirrors the results of Tylén et al., 2015 who found the right hippocampus to be involved in plot formation (Tylén et al., 2015).

However, several semantic memory regions did not show significant synchrony across the three sections. Regions consistently synchronized across all film sections included the left temporal pole, fusiform gyrus, IPS and PFC. The left ACC was synchronized during only the Set-up portion, and the MTG remained synchronous during Set-up and Development but not Resolution. This suggests that semantic memory regions are dynamically engaged and synchronized during narrative progression, but the involvement changes over the course of the narrative. This conclusion is supported by the findings of Jääskeläinen et al., 2020, who found that the synchronous activity in the TPJ, IPS, and FEF are based on the attentional engagement of a narrative. Moreover, a more attentionally engaging film would result in more synchronous activity in these regions. Therefore, changes in synchronous activity in these areas would suggest changes in engagement with narrative relevant content. Additionally, during Set-up, there may be more concerted efforts to establish the narrative’s context and characters, requiring greater synchronous activation within the semantic memory network. As the story progresses, there could be a shift in focus towards more detailed and context-specific processing, leading to reduced synchrony within this network. Thus, while attentional and episodic memory regions remain synchronized across Set-up, Development, and Resolution, neural synchrony in semantic regions (specifically the left ACC and MTG) changes as the narrative progresses.

Together, results from Experiment 1 provide evidence of neural synchrony in the visuospatial attention system (both in the dorsal and ventral attentional networks), the semantic memory system (hubs: temporal pole, ACC, peripheral: MTG, IPS, ACC), and the episodic memory system (precuneus, PCC, AG, and mPFC) for the Set-up, Development, and Resolution phases of a complex audiovisual movie. Further, the study revealed a gradual decline in the engagement of semantic memory areas as the narrative unfolded. This observation is substantiated by the diminishing presence of synchronous activity within semantic regions as the narrative progressed. In the initial Set-up phase, all the regions previously highlighted for their relevance to semantic memory exhibited synchronized activity. However, during the Development portion, the left ACC did not show significant synchronous activity. During the Resolution phase, both the left ACC and MTG exhibited an absence of significant synchronous activity. These patterns of diminishing synchrony suggest a modulation in the cognitive demands associated with semantic memory processing throughout the narrative progression.

## Experiment 2

Narratives undergo constant evolution, continually presenting new information to readers, listeners, or viewers. While Experiment 1 revealed significant neural synchrony within specific sections of the narrative, it did not examine how synchrony differs across the three parts of the narrative. The objective of Experiment 2 was to identify differences in neural synchrony across the three sections of the film, shedding light on the shifting cognitive demands throughout the narrative. Thus, Experiment 2 aimed to identify patterns of neural synchrony for each distinct film section through the following comparisons: Set-up vs. Development, Set-up vs. Resolution, and Development vs. Resolution. The overall hypothesis was that evolving cognitive demands inherent to the narrative would give rise to dynamic alterations in neural synchrony within the attentional, semantic, and episodic networks.

For the Set-up vs. Development contrasts, we hypothesized that regions associated with goal-directed attention would exhibit greater synchrony during the Set-up than the Development phase, semantic memory areas would show greater synchrony during the Set-up phase compared to Development. Conversely, we hypothesized that episodic memory and stimulus-driven visuospatial attention areas would show greater synchrony in the Development than the Set-up phase.For the Set-up vs. Resolution contrasts, we hypothesized that regions involved in goal-directed attention would be more synchronous during Set-up, and regions involved in stimulus-directed attention would be more synchronous during the Resolution phase. Additionally, areas involved in semantic memory would be more synchronous during Set-up than Resolution. Finally, for the Development vs. Resolution contrasts, we hypothesized that regions involved in goal-directed attention would be more synchronous during the Development of the narrative, and regions involved in stimulus-directed attention would be more synchronous during the Resolution phase. Additionally, areas involved in semantic memory would show more synchronous activity during Development than Resolution, while areas involved in episodic memory would be more synchronous during Resolution than Development.

### Methods

The methods for Experiment 2 were the same as Experiment 1, with the following exceptions.

#### Data Analysis

ISC pairings from Experiment 1 were used for this experiment across each film section. To examine differences in paired ISCs, we subtracted the corresponding ISC pairings for each pair of participants for each contrast of interest. Specifically, in the Set-up vs. Development contrast, each of the 190 pairings from the Set-up section was subtracted from its corresponding pair in the Development section using the *fslmaths* ‘-sub’ (Jenkinson et al., 2012) command. This subtraction process was repeated for the Set-up vs. Resolution contrast and the Development vs. Resolution contrast. The resulting subtracted files were analyzed using a one-way LME using the *3dISC* module from AFNI (Chen et al., 2017; Cox, 1996) for each contrast, Set-up > Development, Development > Set-up, Set-up > Resolution, Resolution > Set-up, Development > Resolution, and Resolution > Development, which is equivalent to a series of paired sample *t-* tests. Significant ISC was determined using a voxel-wise FDR threshold of *q* < .05 for each contrast. The critical *t*-value was determined as 4.4804 for Set-up vs. Development, 3.2629 for Set-up vs. Resolution, and 3.5298 for Development vs. Resolution. To account for potential bidirectional synchrony in each contrast, thresholding was performed twice. This means that each contrast could exhibit significant differences in both directions (e.g., Set-up > Development and Development > Set-up for the Set-up vs. Development contrast). Using the critical t-stats, the t-stat image was thresholded using the *fslmaths* ‘-thr’ and ’-uthr’ functions (Jenkinson et al., 2012) at the respective positive and negative critical *t*-values. We then masked the ISC image with the absolute value of the corresponding thresholded *t*-statistic image using the *fslmaths* ‘-mas’ function (Jenkinson et al., 2012).

### Results

#### Set-up vs. Development

##### Set-up > Development

Results from the one-way LME for the Set-up > Development contrast found 11 significant clusters of neural synchrony (see Figure 5 and Supplementary Materials Appendix B), including the right SPL (a dorsal visuospatial attention area), the left precuneus and AG (episodic memory regions), and the left fusiform gyrus and IPS (semantic memory areas). Other regions that showed significant neural synchrony were the left supramarginal gyrus, bilateral precentral gyrus, occipital poles, occipital fusiform gyrus, and left precuneus.

**Figure 5.**
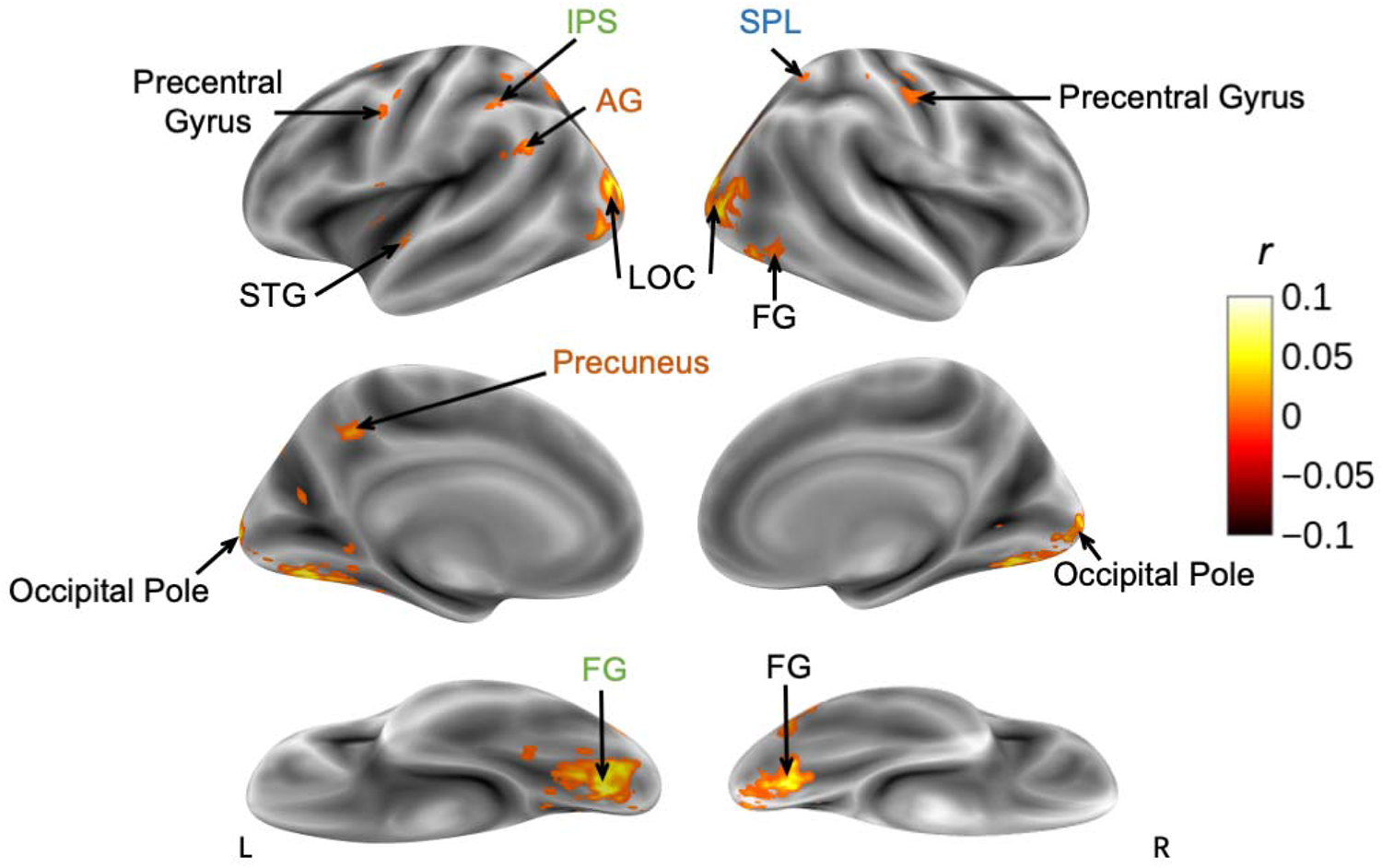
*500 Days of* Summer: Set-up > Development Synchrony. Synchrony maps derived from fMRI data during audiovisual movie watching. Maps are shown for the left (L) and right (R) hemispheres, in the lateral, medial, and inferior orientations. Areas in blue represent neural correlates of visuospatial attention, areas in green represent neural correlates of semantic memory, and areas in orange represent neural correlates of episodic memory.

##### Development > Set-up

Results from the one-way LME for the Development > Set-up contrast found 18 significant clusters of neural synchrony (see Figure 6 and Supplementary Materials Appendix B), including the right IFG/MFG (a visuospatial attention area), and the left AG (an episodic memory area). Other regions that showed significant neural synchrony were the left IFG, right MFG, right temporal pole and areas of the bilateral planum temporale, and left crus II as seen in Figure 6.

**Figure 6.**
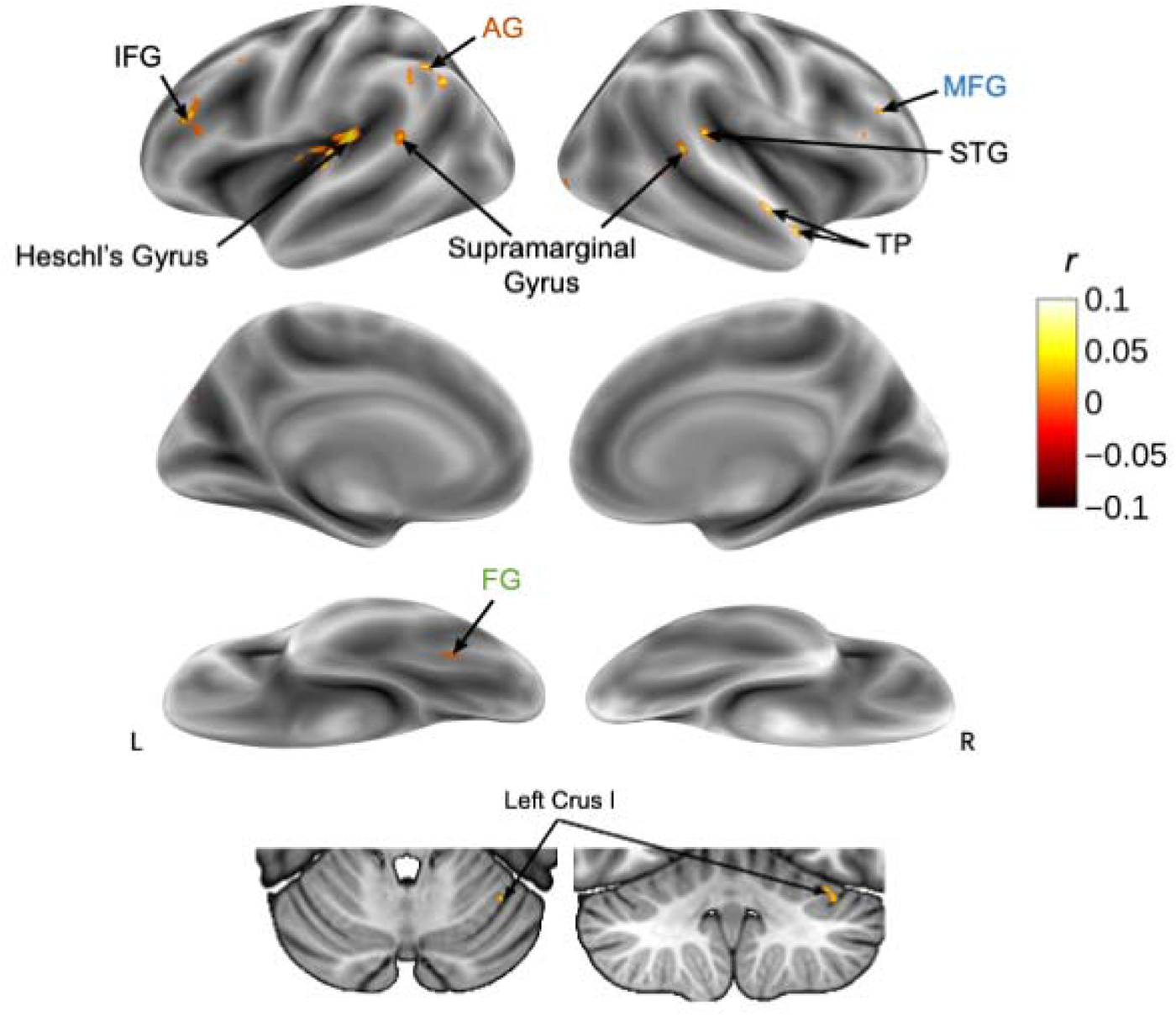
*500 Days of Summer*: Development > Set-up Synchrony. Synchrony maps derived from fMRI data during audiovisual movie watching. Maps are shown for the left (L) and right (R) hemispheres, in the lateral, medial, and inferior orientations. Areas in blue represent neural correlates of visuospatial attention, areas in green represent neural correlates of semantic memory, and areas in orange represent neural correlates of episodic memory.

#### Set-up vs. Resolution

##### Set-up > Resolution

Results from the one-way LME for the Set-up > Resolution contrast identified 43 significant clusters of neural synchrony (see Figure 7 and Supplementary Materials Appendix B), including the right FEF, SPL, and TPJ (visuospatial attention areas), and the bilateral mPFC, precuneus, PHC, RSC, AG, and PCC (episodic memory areas). Semantic memory areas including the hubs of the left temporal pole and ACC, and peripherals including the left fusiform gyrus, MTG, and PFC, also showed significant neural synchrony. Other regions that showed significant neural synchrony were the right thalamus, left crus I, bilateral crus II, bilateral precentral gyrus, left TPJ, and bilateral frontal poles.

**Figure 7.**
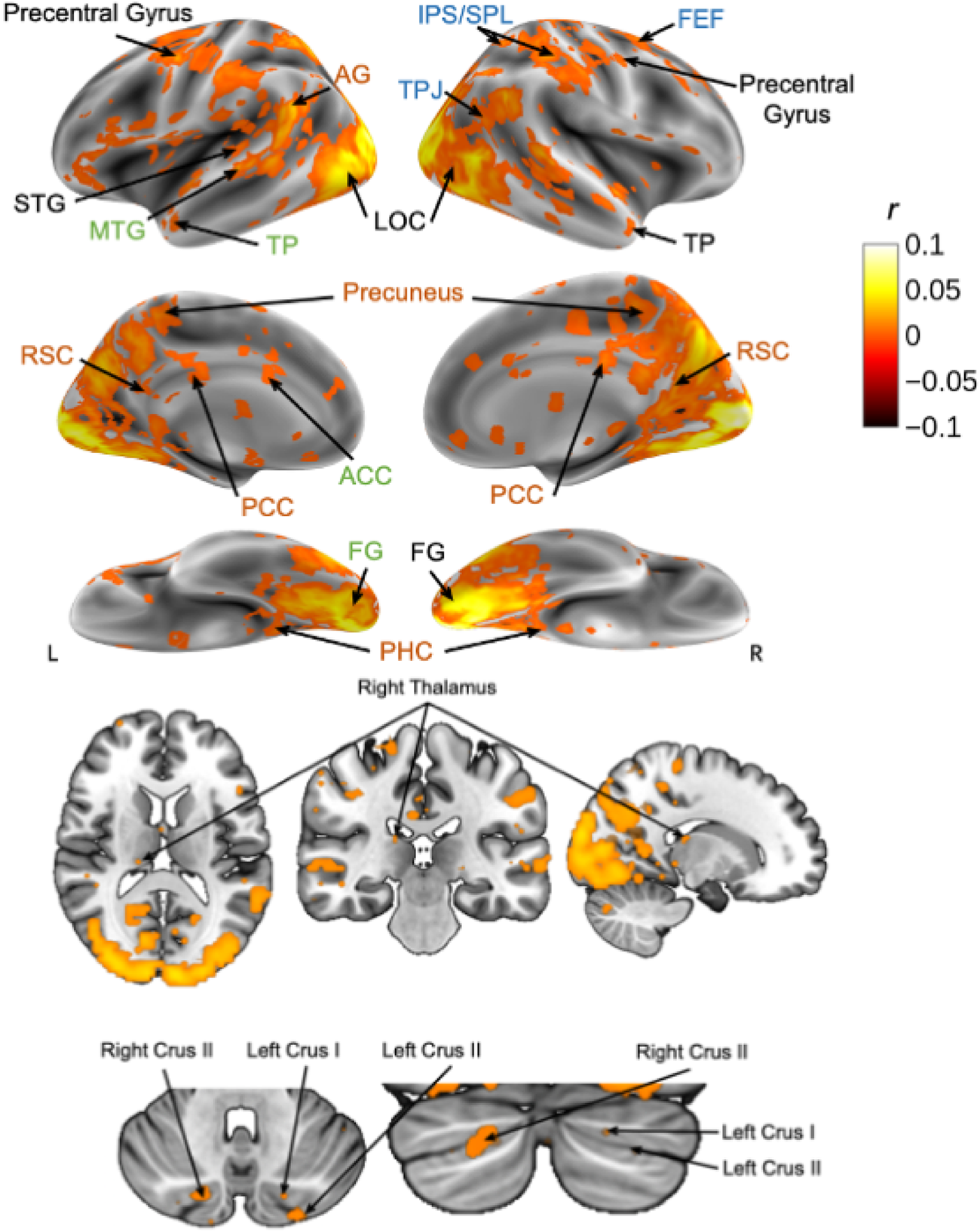
*500 Days of Summer*: Set-up > Resolution Synchrony. Synchrony maps derived from fMRI data during audiovisual movie watching. Maps are shown for the left (L) and right (R) hemispheres, in the lateral, medial, and inferior orientations. Areas in blue represent neural correlates of visuospatial attention, areas in green represent neural correlates of semantic memory, and areas in orange represent neural correlates of episodic memory.

##### Resolution > Set-up

Results from the one-way LME for the Resolution > Set-up contrast identified 14 significant clusters of neural synchrony (see Figure 8 and Supplementary Materials Appendix B). The left ACC, a hub implicated in semantic memory, showed significant neural synchrony. Other regions that showed significant neural synchrony were the right frontal pole, precentral gyrus, and left insular cortex. We did not find significant neural synchrony in visuospatial attention or episodic memory areas.

**Figure 8.**
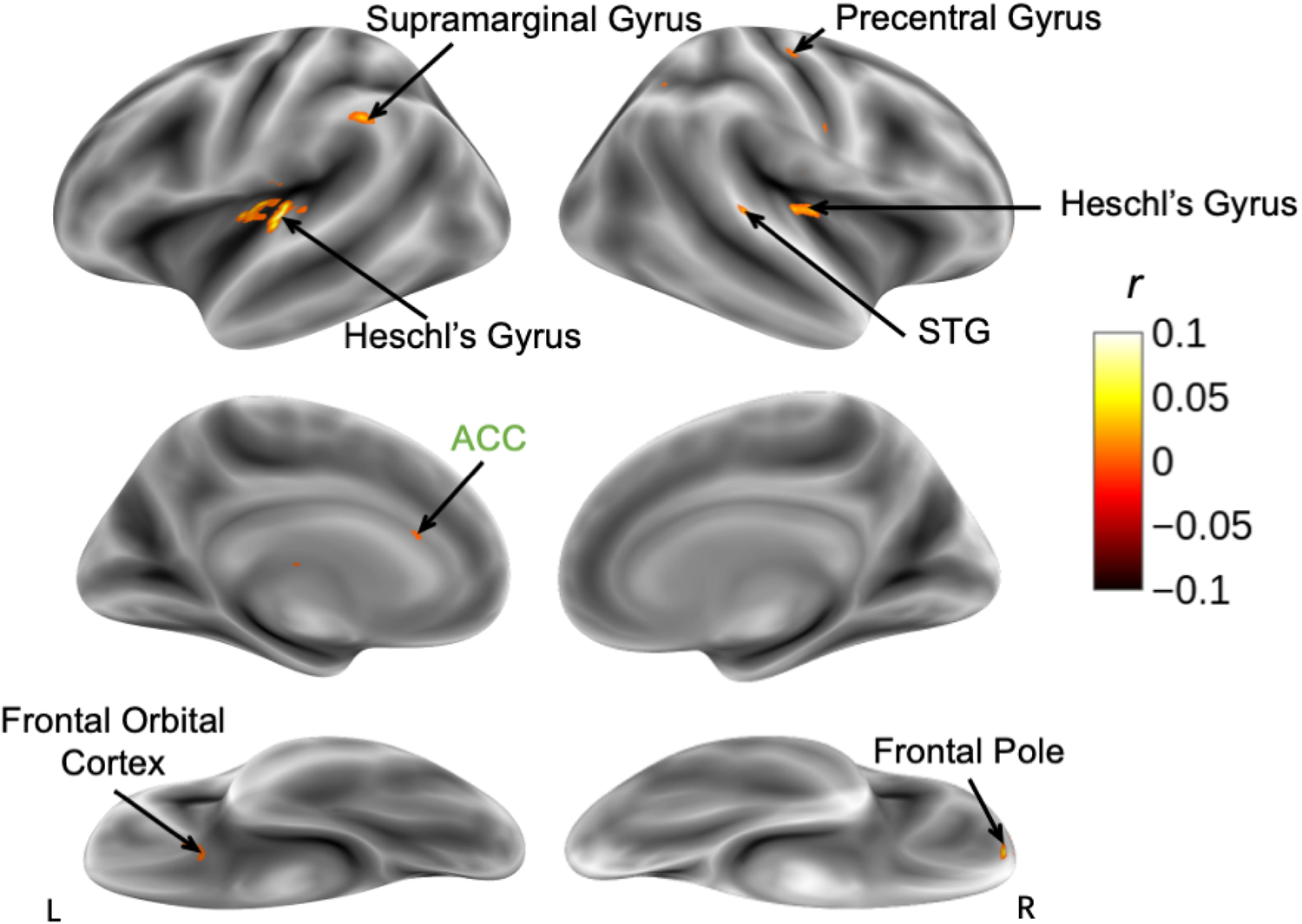
*500 Days of Summer*: Resolution > Set-up Synchrony. Synchrony maps derived from fMRI data during audiovisual movie watching. Maps are shown for the left (L) and right (R) hemispheres, in the lateral, medial, and inferior orientations. Areas in green represent neural correlates of semantic memory.

#### Development vs. Resolution

##### Development > Resolution

Results from the one-way LME for the Development > Resolution contrast found 34 significant clusters of neural synchrony (see Figure 9 and Supplementary Materials Appendix B). Significant clusters in visuospatial attention areas included the right FEF, SPL, TPJ, and IFG/MFG. Significant clusters in episodic memory areas included the left mPFC, bilateral precuneus, PHC, RSC, AG, and PCC. Semantic memory areas, including the hubs of the left temporal pole, and peripherals including the left fusiform gyrus, MTG, and PFC, also showed significant neural synchrony. Other regions that showed significant neural synchrony were the bilateral crus I and left crus II of the cerebellum, the bilateral precentral gyrus, the frontal pole, and left TPJ.

**Figure 9.**
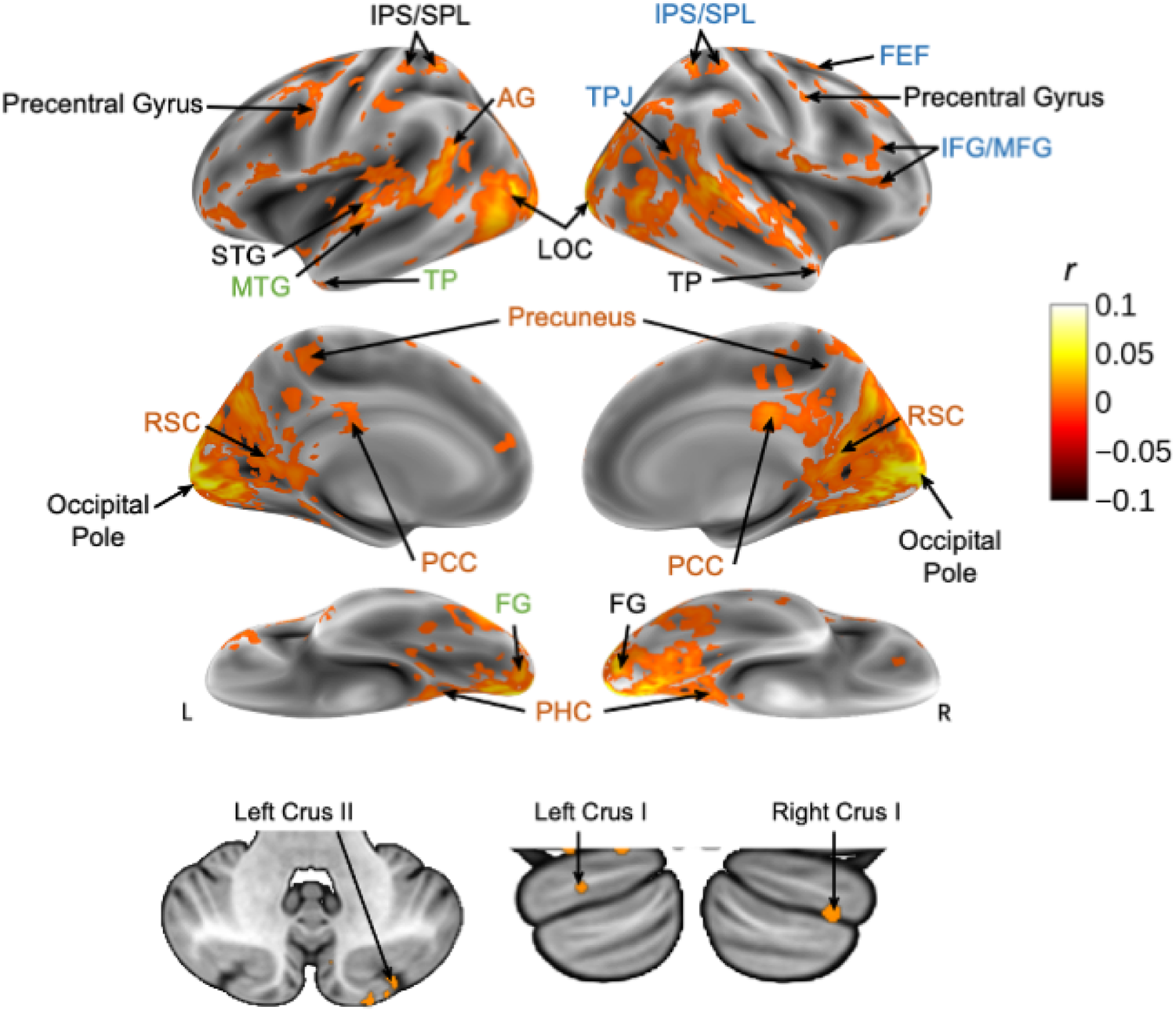
*500 Days of Summer:* Development > Resolution Synchrony. Synchrony maps derived from fMRI data during audiovisual movie watching. Maps are shown for the left (L) and right (R) hemispheres, in the lateral, medial, and inferior orientations. Areas in blue represent neural correlates of visuospatial attention, areas in green represent neural correlates of semantic memory, and areas in orange represent neural correlates of episodic memory.

##### Resolution > Development

Results from the one-way LME for the Resolution > Development contrast did not find any regions with significant synchrony in visuospatial attention, semantic memory, and episodic memory areas. Other regions that showed significant neural synchrony were the right insular cortex, right temporal pole, and inferior temporal gyrus.

**Figure 10.**
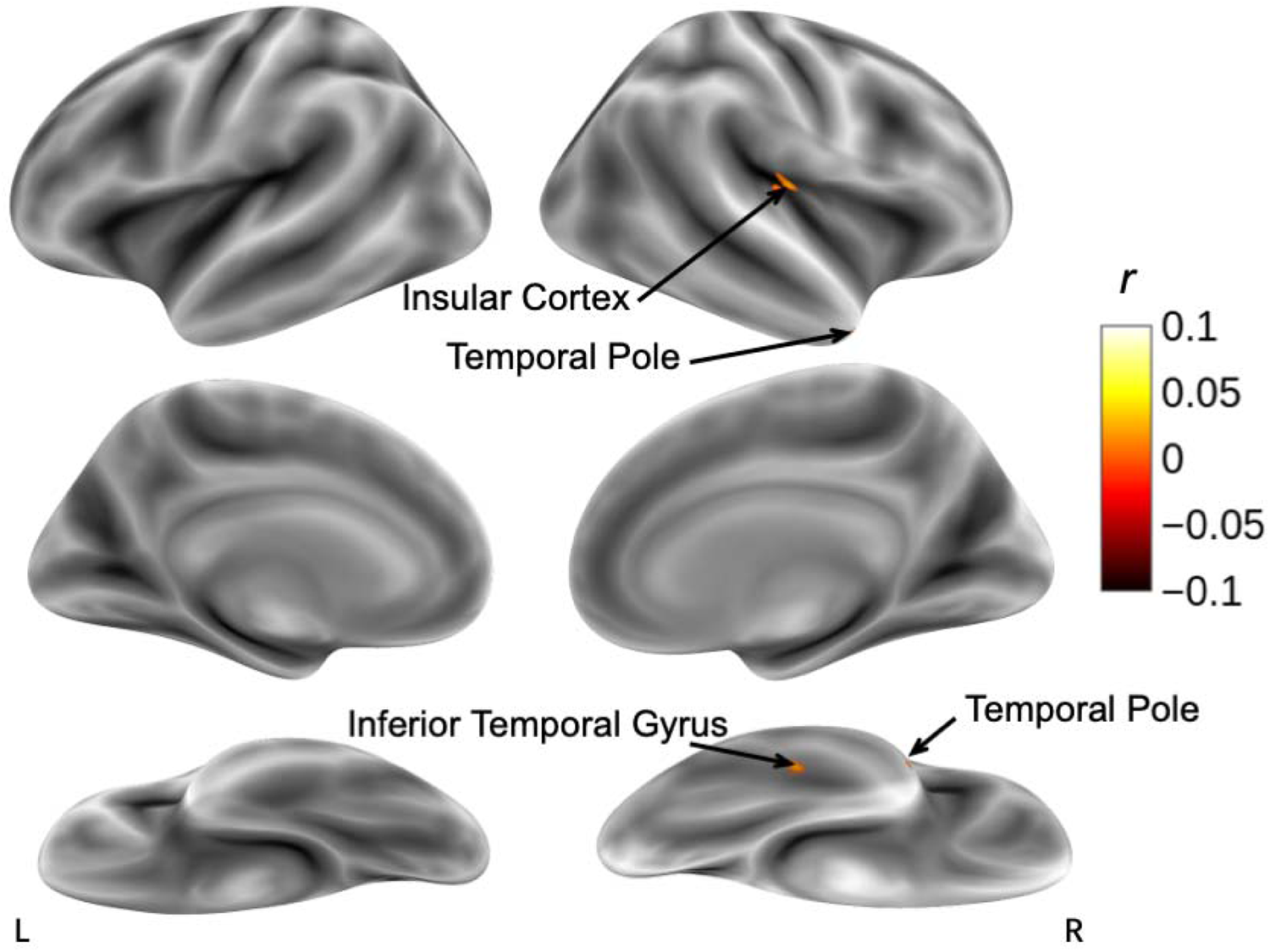
*500 Days of Summer:* Resolution > Development Synchrony. Synchrony maps derived from fMRI data during audiovisual movie watching. Maps are shown for the left (L) and right (R) hemispheres, in the lateral, medial, and inferior orientations.

### Discussion

#### Set-up

The Set-up phase of narrative construction involves establishing the storyline, introducing characters and their connections, and integrating this information. During this phase, there is increased neural synchrony in regions associated with goal-directed attention (right SPL, FEF) and episodic memory (left precuneus, AG). Semantic memory areas (left fusiform gyrus, IPS) also exhibit higher synchrony. This heightened memory engagement aligns with the introduction of characters, their goals, morals, story settings, and relationships. The Set-up phase contrasts with the Resolution phase, showing increased synchrony in attention-related regions (right TPJ) and memory-related areas (bilateral mPFC, PHC, RSC, AG, PCC, left temporal pole, ACC, left fusiform gyrus, MTG, PFC). Other regions with significant synchrony differences include the precentral gyrus (Set-up > Development, Set-up > Resolution), right thalamus (Set-up > Resolution), and bilateral crus II (Set-up > Resolution), suggesting increased demands on discourse comprehension, attention, and social mentalizing during the Set-up phase. Additionally, synchrony in the lateral occipital cortices, occipital poles, and superior temporal gyri suggests processing of audiovisual stimuli.

#### Development

In the Development section, there is increased neural synchrony in regions linked to stimulus-driven attention (right IFG/MFG) and episodic memory (left AG) compared to the Set-up phase. Additionally, attentional regions (right FEF, SPL, IFG/MFG, TPJ) exhibit higher synchrony in Development than in Resolution, suggesting a shift from goal-directed to stimulus-driven attention during this phase. Semantic memory regions (left temporal pole, fusiform gyrus, MTG, PFC) also show heightened synchrony in Development, indicating the integration and retention of new semantic information. These findings reveal evolving cognitive processes during narrative comprehension, with a dynamic shift in attention mechanisms and minimal differences between Set-up and Development. The precentral gyrus’s synchronous activity aligns with its role in discourse comprehension and social cognition, while synchrony in lateral occipital cortices, occipital poles, and superior temporal gyri corresponds to audiovisual stimulus processing.

#### Resolution

When contrasting the Resolution and Set-up phases, patterns of synchrony in the left ACC showed more synchrony during Resolution than Set-up. This region is considered to be a semantic memory hub. When assessing synchrony in semantic and episodic memory, besides the left ACC, there are no other areas within either of these networks that show greater synchrony during Resolution than Set-up or Development. This could suggest that there are significant differences in cognitive demands when comparing the initial stages of a narrative (Set-up and Development) and the end (Resolution), however, visuospatial attention, semantic memory, and episodic memory are less engaged during the final portion of the narrative. This decrease is reflected in the observed lower level of neural synchrony during the Resolution section when contrasted with the Set-up or Development sections.

#### Conclusion

In summary, this experiment offers valuable insight into the dynamic shifts in cognitive demands associated with narrative processing. While previous research has provided evidence of neural correlates connected to narrative processing (e.g., Jääskeläinen et al., 2020; Kauttonen et al., 2018; Tylén et al., 2015), the precise timing of these neural correlates coming into play during the narrative has remained largely unexplored. Here, we examined differences in neural synchrony between the three stages of an evolving narrative. Overall, we found that as the narrative evolves, there are changes in the cognitive demands associated with visuospatial attention, semantic memory, and episodic memory. This finding is supported by the patterns of neural synchrony observed in each of the six contrasts. However, it is important to note that this experiment is focused exclusively on a single narrative following the three-act structure. Consequently, the generalizability of our findings to other narratives may be limited.

Nevertheless, the methodology adopted in this study significantly advances our comprehension of the cognitive processes underpinning narrative processing, which lays the foundation for subsequent experiments.

## General Discussion

We sought to identify the neural correlates of narrative processing in the human brain to elucidate how the communication and coordination between brain regions varied as information was integrated into long-term memory. To do so, we examined how neural synchrony changed over the course of a complex audiovisual narrative. We were specifically interested in how neural synchrony in visuospatial attentional brain regions (i.e., the right FEF, IPS/SPL, IFG/MFG, and TPJ), semantic memory regions (i.e., left temporal pole, ACC, fusiform gyrus, IPS, MTG, PFC), and episodic memory regions (i.e., precuneus, PHC, RSC, AG, mPFC, and PCC) changed over the Set-up, Development, and Resolution sections according to the typical three-act structure of Hollywood films. Our ultimate goal was to provide insight into how narratives become integrated into long-term memory in the human brain using naturalistic stimuli, which better encapsulate the demands of everyday life.

The Set-up phase introduces the main protagonist and sets up the basis of the narrative. Visuospatial attention, semantic memory, and episodic memory areas that showed significant synchrony in Set-up in Experiments 1 and 2 were the right IPS/SPL (involved in goal-directed attention), the left fusiform gyrus (involved in semantic memory), and the bilateral AG and precuneus (involved in episodic memory). Thus, we propose that these regions encompass the core “narrative building” network in the brain and suggest that the initial stages of memory formation rely upon areas involved in voluntary visuospatial attention, episodic memory, and semantic memory, and the integration of this information into coherent narrative frameworks (see Figure 11).

**Figure 11.**
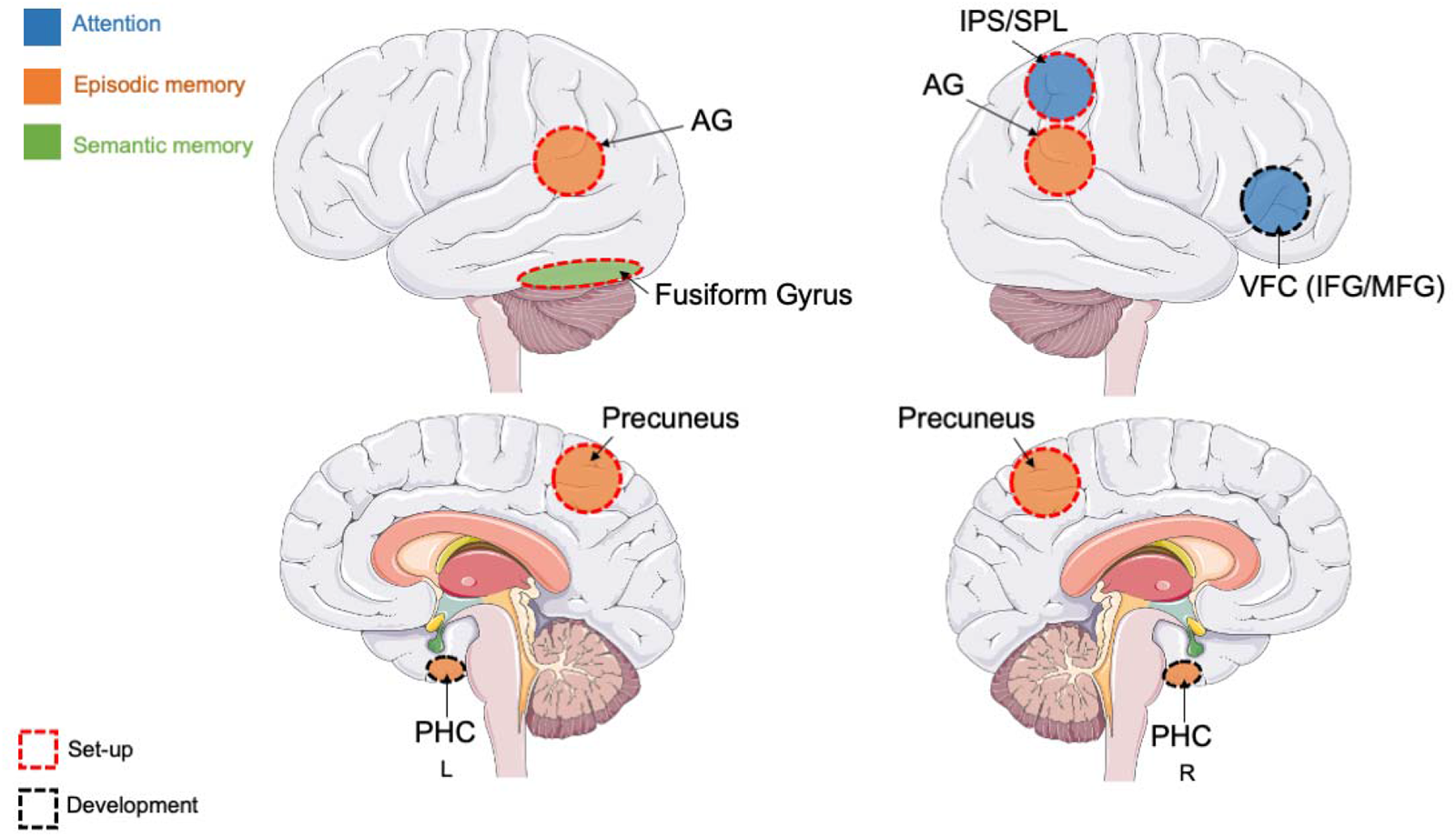
Core Neural Correlates of Narrative Integration: Set-up and Development Core Regions. Illustration of the neural correlates involved in narrative processing. The model proposes that red-outlined regions are associated with the Set-up phase and the black outline is implicated in the Development phase. The model provides a comprehensive framework for understanding the neural correlates involved in processing narratives over extended timescales. Brain model downloaded from Servier Medical Art (“SMART – Servier Medical ART,” n.d.) licensed under a Creative Commons Attribution 3.0 unported license.

Koenigs et al. (2009) found that damage to the SPL resulted in significant deficits on tests involving the manipulation and updating of working memory information. This suggests the SPL may play an important role in the construction and maintenance of narrative representations. The AG, a region implicated in episodic memory, is involved in multimodal sensory integration during both encoding and construction of coherent episodic memories (Tibon et al., 2019) and enable vivid recall during both encoding and retrieval (Ritchey & Cooper, 2020; Tibon et al., 2019). Furthermore, the precuneus is involved in updating mental story representations during narrative processing (Whitney et al., 2009) and narrative interpretation (Jääskeläinen et al., 2020). Our study, in addition to the above literature, implicates the SPL, AG, and precuneus in episodic memory processes crucial to narrative processing.

### The Neural Correlates of Developing a Narrative: Development

During the Development phase, the protagonist encounters progressively difficult obstacles as the narrative unfolds, they confront the most pivotal obstacle, intensifying tension and driving the narrative toward its climax. Based on the results of Experiments 1 and 2, we identified a core narrative development network that includes the right IFG/MFG and the bilateral PHC. This is a particularly interesting finding, as it suggests that these areas may be involved in integrating information into long-term memory over longer timescales than the regions found in the Set-up portion. In contrast to Set-up, involvement of the right IFG/MFG later in the narrative may be associated with a shift from voluntary attention to stimulus-driven attention (see Figure 11). Previous research examining narrative processing in individuals with unilateral brain injury found that damage to the IFG/MFG resulted in deficits in processing more complex narratives (Karaduman et al., 2017). Additionally, the PHC is implicated in memory system networks as a critical support of memory processes (Ward et al., 2014). Additionally, the PHC are critically affected in the manifestation of Alzheimer’s disease (Jacobs et al., 2012; Wang et al., 2012). These findings, in conjunction with our results, suggest the critical role of the PHC in the building and updating of information during long-term episodic memory processes within narrative processing.

### The Neural Correlates of Concluding a Narrative: Resolution

In the Resolution, the protagonist overcomes the most significant challenge, leading to the climactic moment and eventual resolution. The results of Experiment 1 and 2 show no areas within the visuospatial attention, semantic memory, or episodic memory networks consistently synchronous across all 3 contrasts (Experiment 1: Resolution, Experiment 2: Resolution > Set-up and Resolution > Development). The inconsistency in synchrony observed in Resolution, especially when compared to Set-up and Development, indicates significant variations in cognitive demands associated with processing narratives. The initial stages (Set-up and Development) appear to play a crucial role in creating and sustaining coherent narrative representations.

Together, this research provides evidence of evolving cognitive demands associated with the 3-act narrative structure results in dynamic changes in neural synchrony, specifically in terms of attention, semantic memory, and episodic memory. We identified two core narrative processing networks involved in building coherent representations over extended timescales. The neural correlates of these networks include the right IPS/SPL, IFG/MFG (visuospatial attention), bilateral, precuneus, AG, PHC (episodic memory), left fusiform gyrus (semantic memory). These regions may form the scaffolding for successful narrative processing over the course of naturalistic stimuli. This work offers valuable insight into narrative processing on extended timescales, mirroring processing that is more applicable to “real-world” scenarios and establishes a framework of neural correlates of long-term memory formation.

The current study extends research focused on examining neural activity associated with naturalistic stimuli. Our novel approach of examining changes in neural synchrony associated with different stages of narrative processing (i.e., Set-up, Development, and Resolution) offers valuable insight into not only narrative processing as a whole, but also dynamic changes in cognitive demands. Previous research has generally employed brief naturalistic stimuli (Kauttonen et al., 2018; Tylén et al., 2015; Xu et al., 2005), limiting the ability to examine changing neural correlates over extended timescales. In contrast, the experiments in this study investigated a larger spectrum of narrative evolution, encompassing the unfolding cognitive engagement from inception to resolution within the three-act structure. Our results also have important implications for understanding disorders marked by the breakdown of narrative processing, which may in turn help advance both clinical interventions and theoretical frameworks in the realm of cognitive neuroscience.

### Limitations and Future Directions

This study aimed at uncovering the neural underpinnings of narrative processing, however, there are several limitations to be discussed. Because we used data from an online database, we did not have control over the stimuli that were presented to the participants, limiting our control over plot content, length, and genre of the films chosen. In addition, the data used in these experiments was collected using a 1.5T MRI, which has lower spatial specificity and a lower signal-to-noise ratio than stronger MRIs. The use of a higher Tesla scanner would better encapsulate finer details of functional activations. Future research should aim at using higher-resolution MRI instruments to acquire such data.

### Conclusion

In conclusion, this study explored narrative processing across three distinct phases of complex audiovisual narratives using naturalistic stimuli. We identified neural correlates associated with visuospatial attention, semantic memory, and episodic memory within the Set-up, Development, and Resolution segments of the narrative. The findings collectively emphasize dynamic changes in neural synchrony associated with evolving cognitive demands intrinsic to narrative structure. We found that the initial portion of narrative processing (Set-up) relies upon areas involved in goal-directed visuospatial attention (right IPS/SPL), episodic memory (bilateral AG, precuneus), and semantic memory (left fusiform gyrus). Areas significantly involved in Development include areas involved in stimulus-driven visuospatial attention (right IFG/MFG) and episodic memory (bilateral PHC). Our results also infer that these areas are involved in the formation of narrative representations that allow us to integrate information over extended timescales. In summary, the research presented in this study provides valuable insight into how knowledge is encoded, consolidated, and retrieved in the brain using naturalistic stimuli, shedding light on the fundamental mechanisms that shape cognition. This may inform strategies for enhancing memory-related interventions and treatments.

