## Supplementary Materials for "Mapping the Changing Neural Architecture of Narrative Processing Using Naturalistic Stimuli: an fMRI Study"

Supporting Information Appendices for a manuscript: Haines C., Craig J., Klamer K., Sullivan K., Ekstrand C.: “Neural Correlates of Narrative Structure During Naturalistic Audiovisual Film Using Functional Magnetic Resonance Imaging

**List of Appendices:**

A: Tables for Experiment 1

B: Tables for Experiment 2

A. **Tables for Experiment 1**

Table 2.1. *500 Days of Summer*: Set-up Cluster Table. Coordinates in MNI space. Only clusters with five or more voxels are reported. Voxels are visualized in Figure 2.

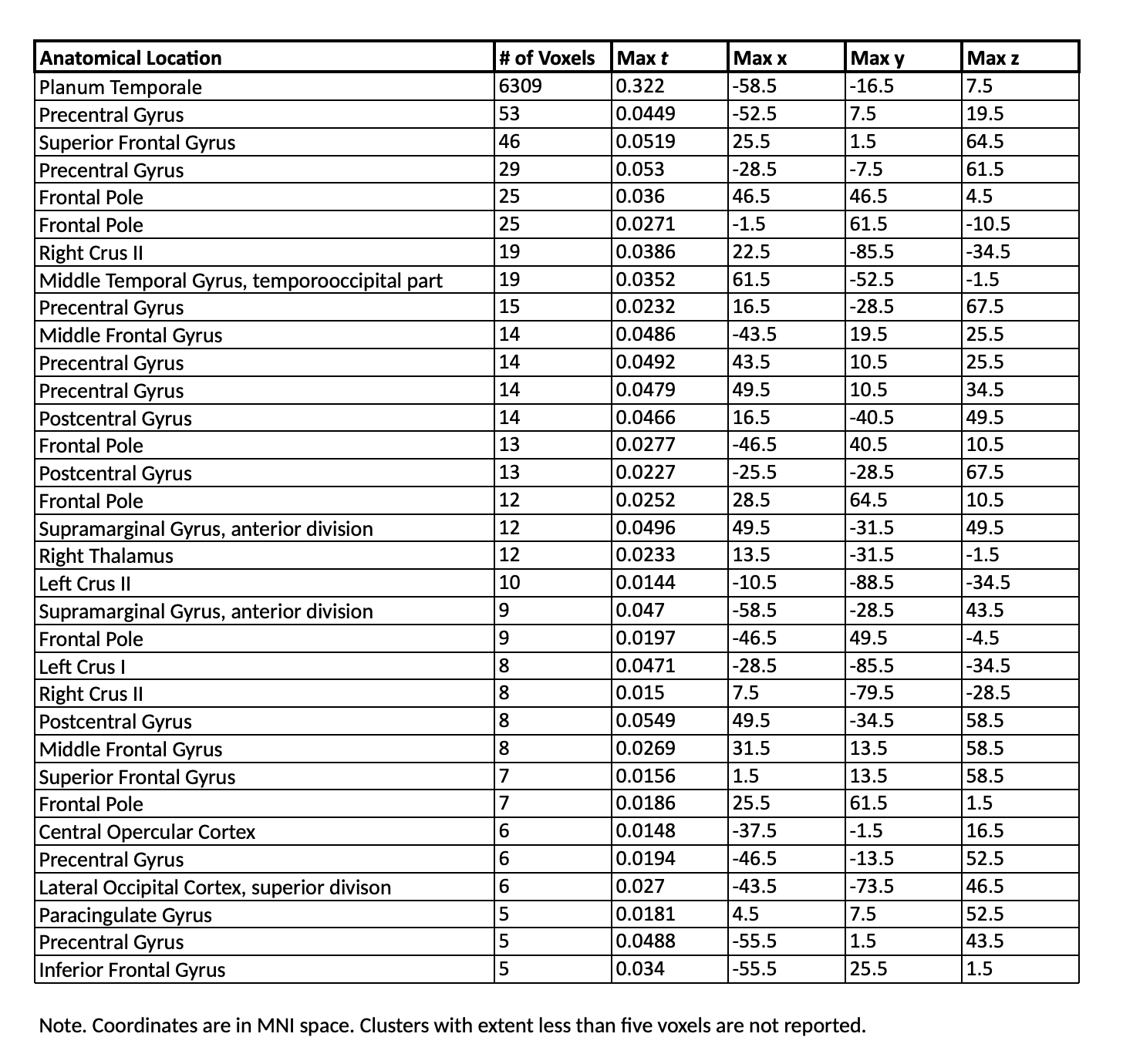

Table 2.2. *500 Days of* Summer: Development Cluster Table. Coordinates in MNI space. Only clusters with five or more voxels are reported. Voxels are visualized in Figure 3.

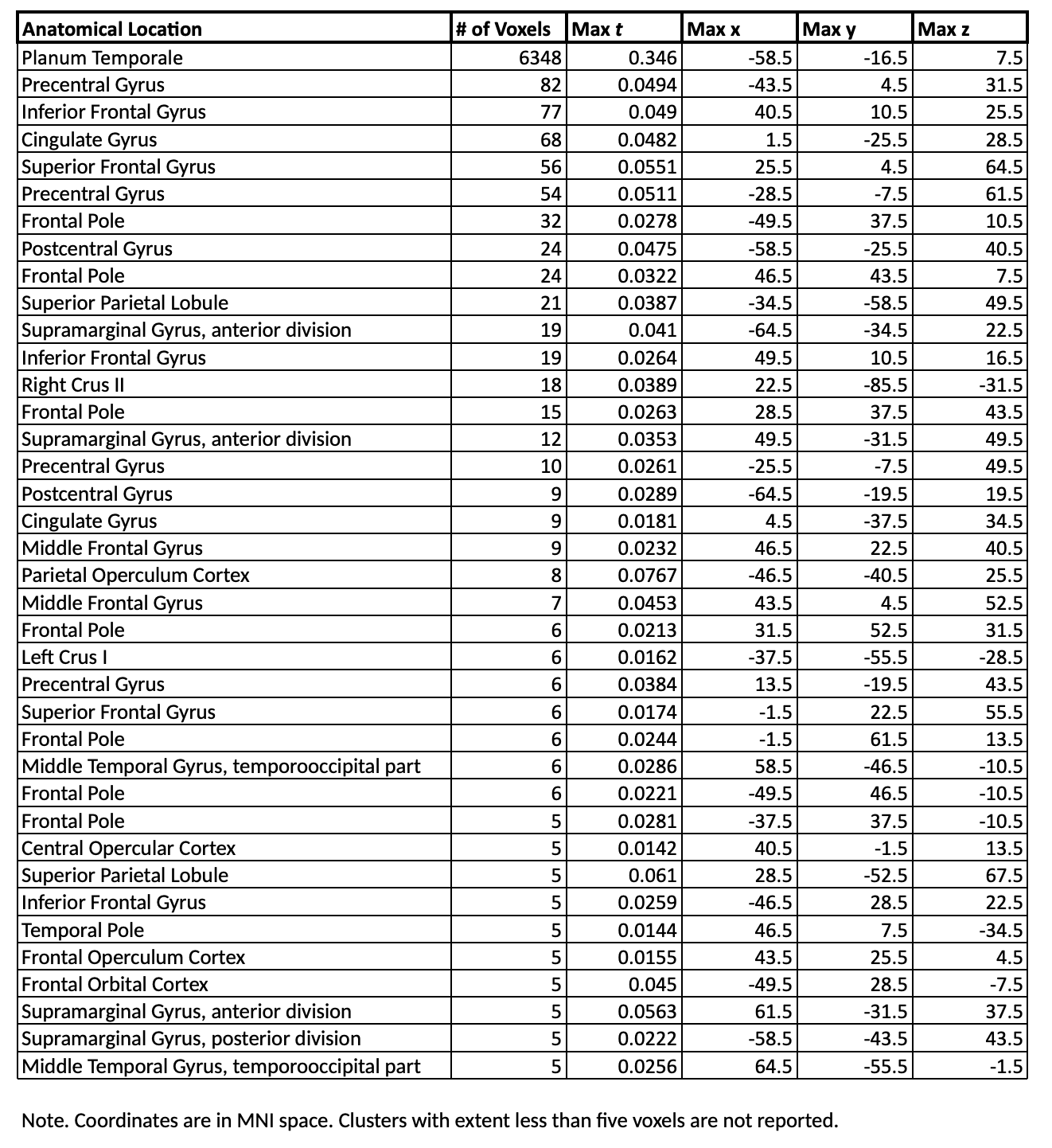

Table 2.3. *500 Days of* Summer: Resolution Cluster Table. Coordinates in MNI space. Only clusters with five or more voxels are reported. Voxels are visualized in Figure 4.

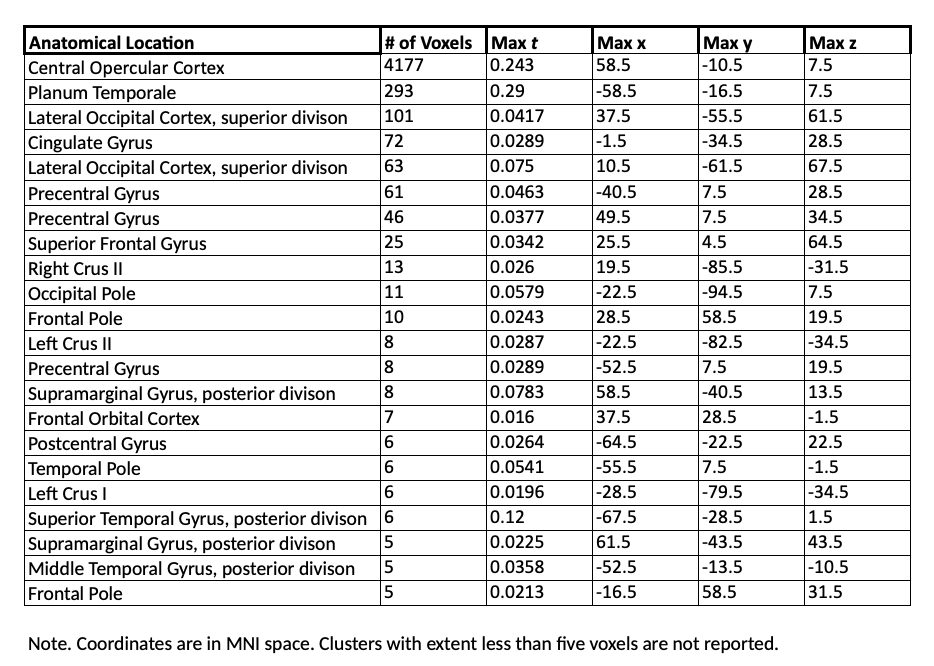

B. **Tables for Experiment 2**

Table 2.4. *500 Days of* Summer: Set-up > Development Cluster Table. Coordinates in MNI space. Only clusters with five or more voxels are reported. Voxels are visualized in Figure 5.

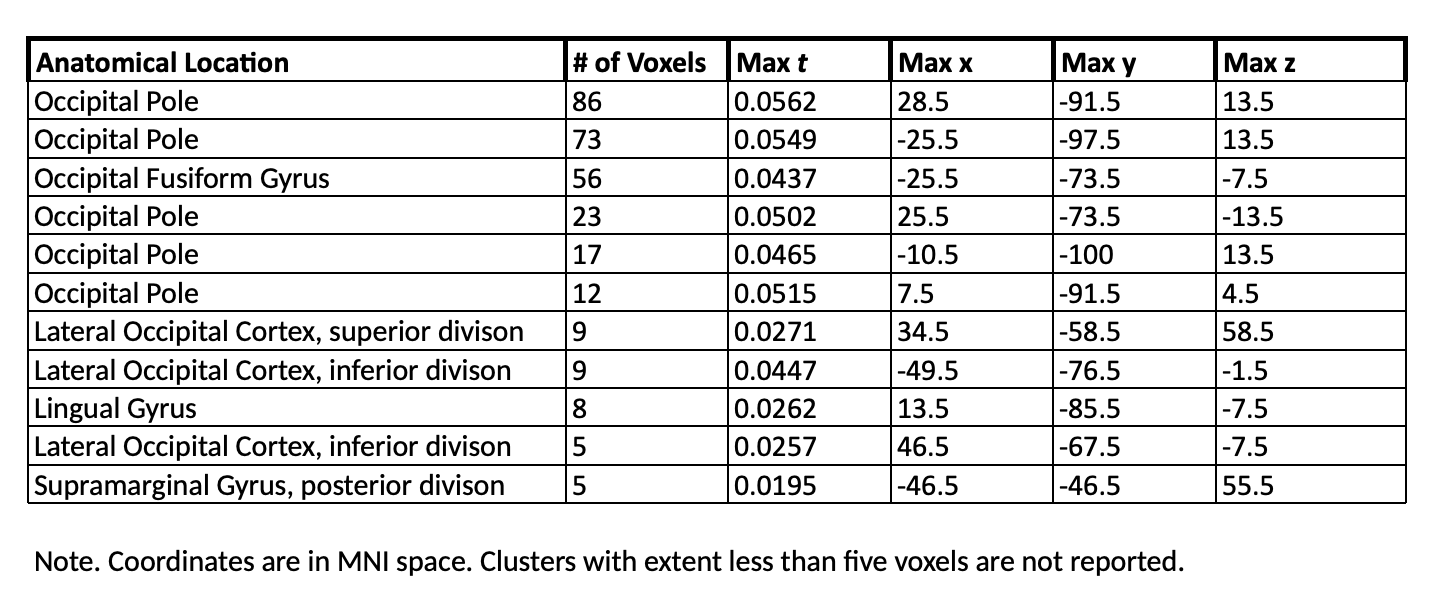

Table 2.5. *500 Days of Summer:* Development > Set-up Cluster Table. Coordinates in MNI space. Voxels are visualized in Figure 6.

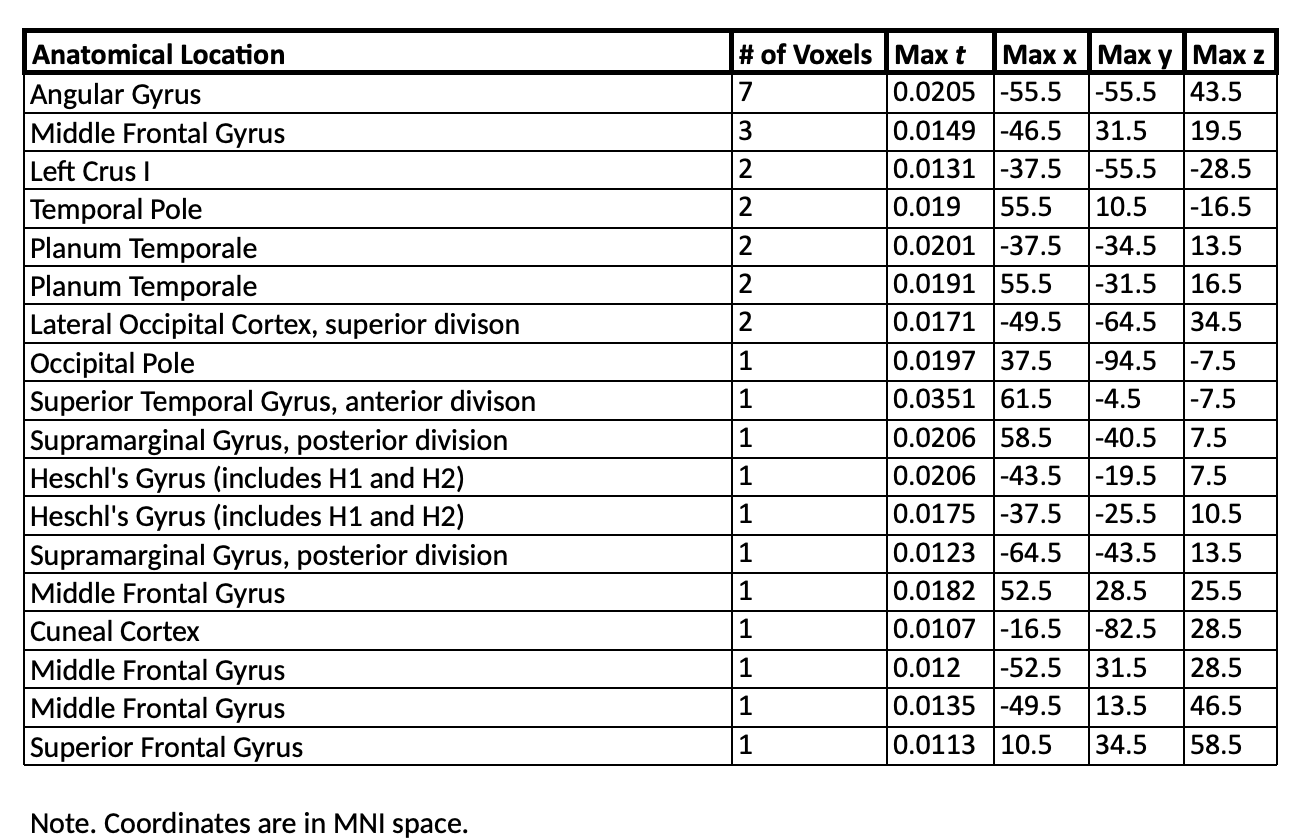

Table 2.6. *500 Days of Summer:* Set-up > Resolution Cluster Table. Coordinates in MNI space. Only clusters with five or more voxels are reported. Voxels are visualized in Figure 7.

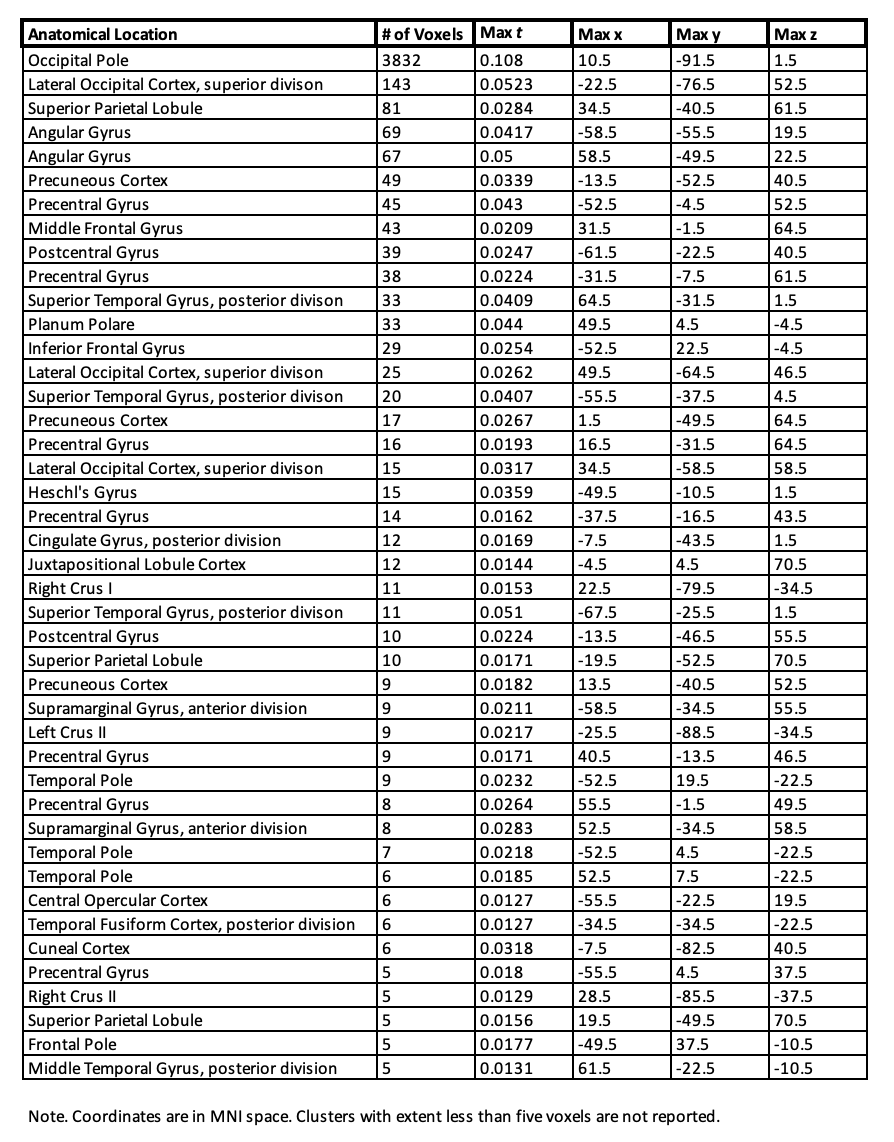

Table 2.7. *500 Days of Summer:* Resolution > Set-up Cluster Table. Coordinates in MNI space. Voxels are visualized in Figure 8.

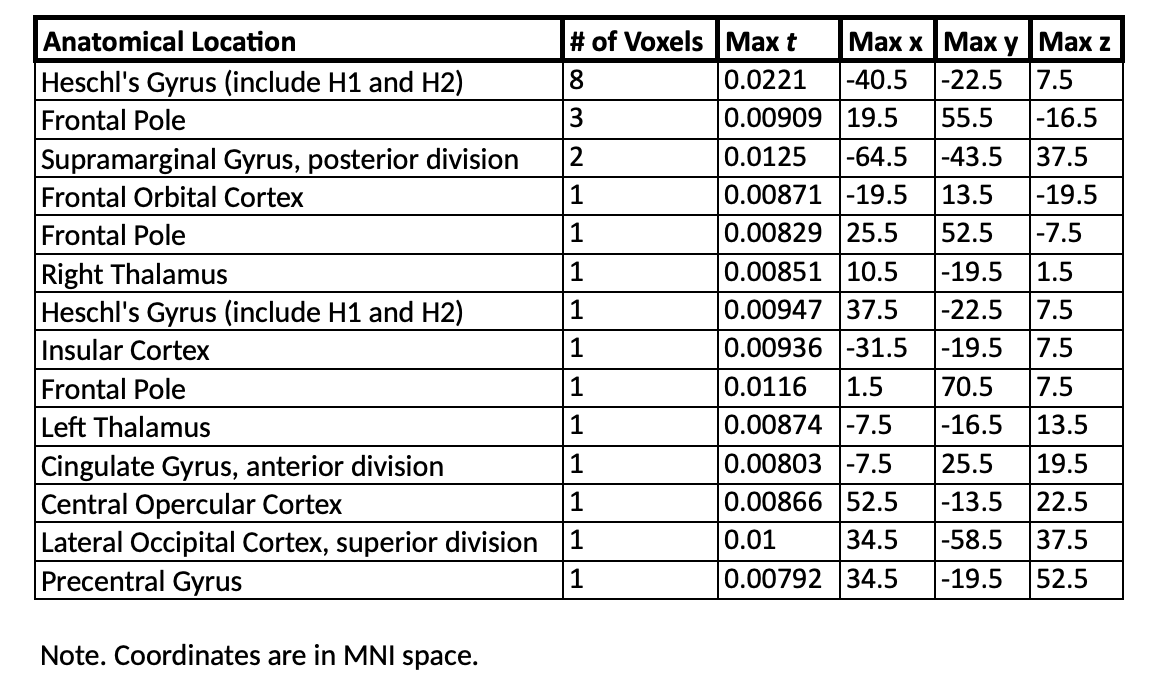

Table 2.8. *500 Days of Summer:* Development > Resolution Cluster Table. Coordinates in MNI space. Only clusters with five or more voxels are reported. Voxels are visualized in Figure 9.

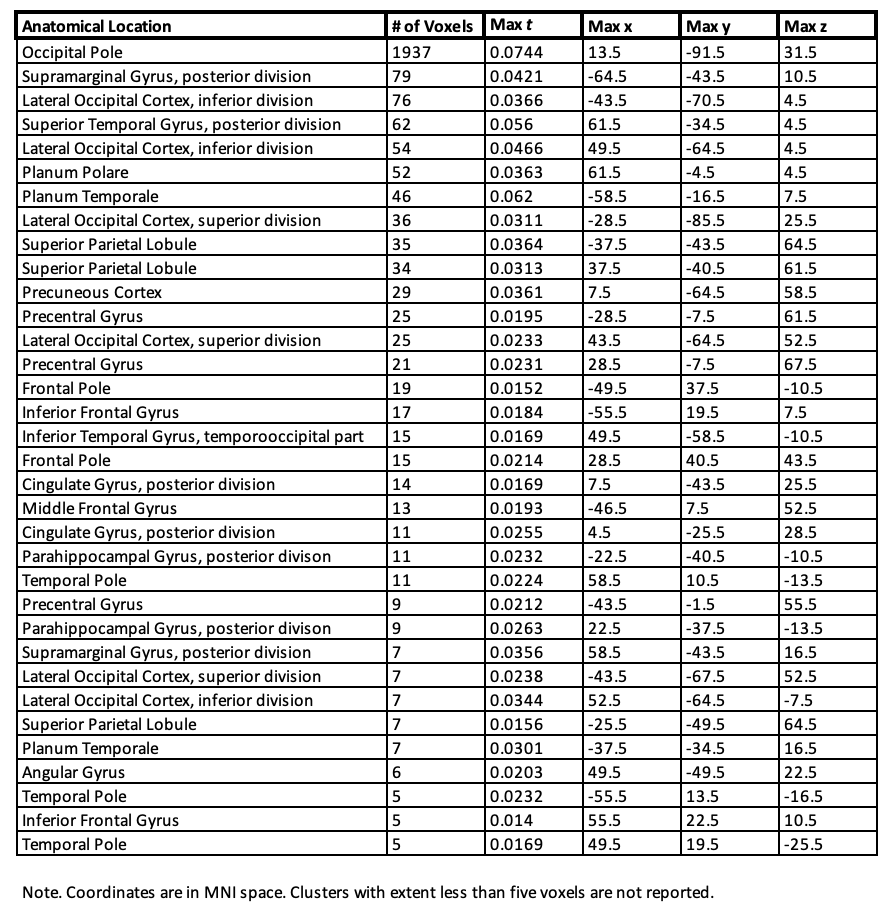

Table 2.9. *500 Days of Summer:* Resolution > Development Cluster Table. Coordinates in MNI space. Voxels are visualized in Figure 10.

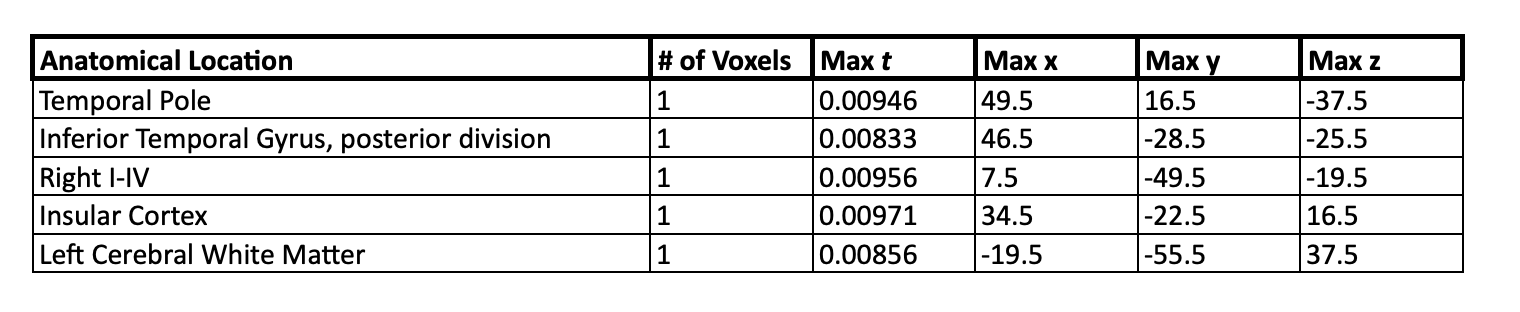
